# Understanding the role of chromatin accessibility and compaction on loop extrusion dynamics

**DOI:** 10.64898/2026.10.12.758711

**Authors:** Nicolas Pellet, César L. Pastrana, Axel Delamarre, Dario D’Asaro, Pascal Bernard, Daniel Jost

## Abstract

Loop extrusion by SMC complexes is a fundamental mechanism regulating the three-dimensional genome organization, but how these molecular motors navigate through the crowded, nucleosome-covered chromatin fiber remains an open question. Here, we develop a stochastic biophysical model of loop extrusion that explicitly couples motor stepping dynamics to chromatin accessibility and compaction. First, we systematically analyze the role of accessibility on its dynamics across homogeneous, periodic, and random roadblock positioning along chromatin. This comprehensive analysis reveals distinct dynamic regimes: phase-dependent kinetics in periodic arrays where motor step size aligns with obstacle spacing, and anomalous sub-linear progression in disordered environments due to rare but long-lived trapping events. Then, we contextualize our framework to nucleosome-like arrays. Our simulations reveal a non-intuitive dual role for nucleosomes: while they hinder extruder progression by blocking access to linker DNA, their high compaction packs more base pairs per physical distance, enabling larger genomic displacements per step. Strikingly, the resulting base-pair extrusion speed displays a non-monotonic dependence on nucleosome spacing, reaching a maximum at intermediate linker lengths (10-70 bp) that can exceed the speed on naked DNA. Finally, in support of this mechanism, analysis of yeast genomic data shows that a higher nucleosome occupancy between cohesin-associated regions correlates with stronger contact enrichment, consistent with compaction-enhanced extrusion efficiency in vivo. These findings establish chromatin architecture as a tunable regulator of loop extrusion, explaining the comparable speeds observed in vitro and in vivo and providing a quantitative framework for understanding how genome organization emerges from the interplay between motor kinetics and chromatin structure.

## 1. Introduction

The three-dimensional organization of the genome within the nucleus of eukaryotic cells constitutes a distinct level of biological regulation, separate from the DNA sequence. [1, 2]. Far from being randomly distributed throughout the nuclear volume, chromatin is organized according to a hierarchy of structured functional scales [3–11]. At large scales (1−10*µm*, 1 −100Mbp), each chromosome occupies a relatively delimited territory and the genome segregates into two compartments, A and B, corresponding respectively to transcriptionally active and repressed regions. At an intermediate scale (100s nm, 100s kbp), topologically associating domains (TADs) are contiguous regions of the genome where DNA sequences interact with one another in three-dimensional space more frequently than with sequences outside the domain. These domains are often marked by specific boundary elements—such as CTCF-binding sites in mammals—and are believed to organize genes together with their regulatory sequences [12]. Finally, at a smaller scale (10 − 100nm, 1 −100kbp), specific loops bring distant regulatory elements, such as promoters and enhancers or TAD boundaries, into physical proximity [13, 14].

One key molecular mechanism involved in the emergence of this structural hierarchy is active DNA loop extrusion by SMC (Structural Maintenance of Chromosomes) complexes [15–18]. In this process, loop extruding factors (LEF), such as cohesin or condensin, load onto the chromatin fiber and translocate along the DNA, progressively enlarging a loop until they encounter barriers (eg, CTCF-binding sites for cohesin) or until they dissociate [16, 18, 19]. This model, formalized about a decade ago, naturally predicts the formation of TADs displaying the characteristic corner loop patterns seen in Hi-C maps and the dependence of domain boundaries on the orientation of CTCF motifs [12, 18]. The predictive power of the loop extrusion mechanism has since been confirmed by several depletion experiments of LEF subunits or cofactors [12, 20–22], even though alternative or complementary SMC-driven mechanisms continue to be debated within the community [23, 24]. In addition of this indirect in vivo evidence, the loop extrusion process has received strong in vitro direct experimental support. Real-time visualization of extrusion, obtained by single-molecule imaging on bare DNA stretched by hydrodynamic flow, revealed the three eukaryotic SMC complexes (cohesin, condensin and SMC5/6) capable of generating a growing loop in the presence of ATP, at a speed on the order of 0.5 to 2 kb/s and with low stalling force [25–29]. Magnetic tweezer experiments subsequently showed that extrusion proceeds through discrete steps [30–33]. These results confirm the general character of this mechanism among SMC complexes, regardless of the exact molecular behavior [34–37].

However, these in vitro experimental foundations were established in radically simplified environments in comparison to the complex state of the cell nucleus: bare DNA, in dilute solution, devoid of the many bound proteins, including nucleosomes, that normally populate chromatin in vivo. Experiments exploring the interplay between extruder progression and the presence of roadblocks, reveal a complex crosstalk. Isolated nucleosomes, RNA polymerase, DNA-bound proteins, or even artificial nanoparticles far exceeding the size of the SMC ring offer, individually, only minimal resistance to the progression of condensin or cohesin [38] — a permissiveness consistent with the observation that extruding condensin complexes can themselves bypass one another [39]. In contrast, dense and repeated arrays of DNA-bound proteins can completely block the progression of the extruder. Indeed, Analikwu et al. demonstrated in vitro [40] that regular and dense coverage of telomeric-like DNA by the Rap1 protein stalls the extrusion of yeast condensin, the stalling efficiency depending directly on the length of the protein array and on the density of occupied binding sites. These results highlight a fundamental distinction between isolated and repeated obstacles: the very same type of protein-DNA interaction can shift from negligible resistance to complete blockage depending on the roadblock density and spatial organization along the DNA template — a nuance that considerably complicates the picture of an extruder able to indifferently traverse its nuclear environment, and one that echoes the more general observation that condensin preferentially associates with nucleosome-depleted genomic DNA in vitro [41, 42] and in vivo [43].

Reconciling these observations with the cellular context therefore requires accounting for the nucleosome itself as the most pervasive obstacle that an extruder can encounter. In vivo, nucleosome positioning is known to play a major role in shaping genome architecture at multiple scales [44–46]. However, its more local function of acting as a discrete physical obstacle and constraining the accessibility of the underlying DNA to extrusion remains comparatively poorly understood [47–51]. Knowing the large variety and heterogeneity of nucleosomal landscapes observed within and across species from quasi-periodic to fuzzy patterns [46], this raises a broad open question: how loop extrusion behaves in a crowded, heterogeneous, and complex chromatin environment. In this work, we address this question using biophysical modeling.

In the past decade, polymer and biophysical models have made significant progress in predicting global emerging consequences of extrusion, reproducing loop and domain formation [52–59], and estimating extruder processivity from experimental data [14, 60–62]. However, only very few explored the impact of roadblocks on the loop extrusion dynamics. Using stochastic simulations of SMC diffusion on unidimensional arrays in the presence of obstacles, Maji et al. [63] suggest that, beyond a certain density, the interplay between nucleosome and linker sizes ultimately accelerates cohesin diffusion along chromatin, as the difficult bypass of an obstacle can allow a greater number of base pairs to be traversed. In contrast, recent molecular dynamics simulations of condensin loop extrusion on obstacles-bound DNA arrays by Dooms et al. [64] show that, if sufficiently tightly packed, obstacles impede the ability of the extruder to capture long DNA steps.

In the present study, we address the dual role of nucleosomes on loop extrusion progression: as obstacles limiting the local accessibility to free – extrudable – linker DNA and as compact objects encompassing more base-pairs per unit length than linker DNA. Using stochastic simulations and analytical calculations of a simple biophysical model of unidirectional loop extrusion, we demonstrate that the arrangement and density of roadblocks play an essential role in regulating loop extrusion dynamics and that the higher compaction of DNA around nucleosomes in vivo may actually compensate for the restricted accessibility to linker DNA and may lead to DNA extrusion speed higher than on bare DNA. We confront our model predictions with experimental data in yeast, suggesting that the compaction effect of nucleosomes may be dominant in vivo.

## II. Materials and Methods

### A. Model

We develop a phenomenological model for one-sided loop-extruding factors (LEFs) progressing stochastically along a chromatin fiber where some regions can be more or less accessible to LEF. Our framework couples the motor stepping activity of LEFs [13, 25, 27–32, 65] to the heterogeneous accessibility and compaction of chromatin (Fig.1) and quantifies how this interplay may control loop extrusion dynamics.

**FIG. 1.**
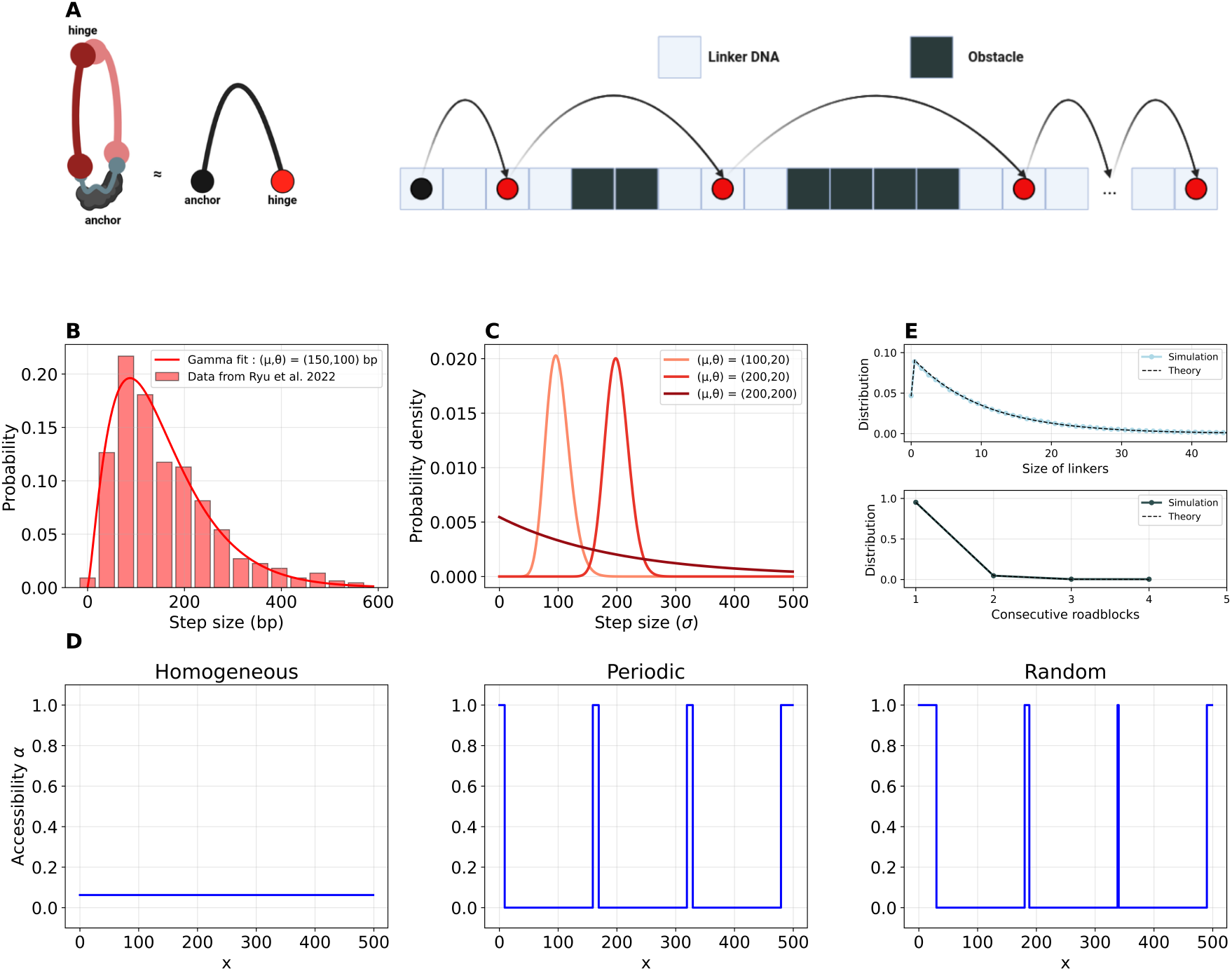
Model of one-sided loop extrusion. **(A)** Schemes of one sided loop extruder such as condensin (Left) and of our model of loop extrusion (Right) with stochastic jumps along a 1D lattice in presence of obstacles (nucleosomes). **(B)** In vitro behavior of a one-sided loop extrusion factor (condensin) measured using magnetic tweezers under a force of 0.4 pN and fitted with a Gamma distribution (adapted from [31]). **(C)** Gamma Law expressed by different mean values *µ* and different standard deviations *θ*. **(D)** The three different landscapes generated for *s* = 150 and 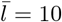 *homogeneous* (left) with a density 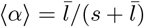, *periodic* (center) and *random* (right). **(E)** Distribution of linker sizes (top) and of the number of consecutive roadblocks (bottom). Simulation fits theory.

The chromatin fiber is modeled as a 1D lattice of size *L*_max_ indexed by position *x* (Fig.1 A, Right). Each site *x* corresponds to a fixed length *σ* of chromatin (given in *nm*), comprising a number of base-pairs (bp) *c*(*x*) that may vary depending on the position and on the local compaction of the chromatin. For each site, we define an accessibility variable *α*(*x*) that reflects the ability of the LEF hinge to bind to it.

The one-sided loop extrusion on a chromatin fiber is modeled as the step-wise stochastic progression of the LEF hinge from left to right along the lattice, the LEF head being assumed immobile at position *x*_0_ (Fig.1A). The hinge, initially at position *x*_0_, attempts to move forward at fixed rate *k*_0_ by “jumping” from sites to sites. At a given time *t* and position *x*, the hinge tries to capture a chromatin segment of size Δ*x* to enlarge the extruded loop, randomly drawn from a fixed distribution *p*(Δ*x*), and this trial move is accepted with a probability *α*(*x* + Δ*x*). This progression is simulated with the Gillespie algorithm [66, 67] with position-dependent propensities given by {*k*_0_*p*(Δ*x*)*α*(*x* + Δ*x*), for Δ*x* ≥ 1}. Each trajectory is thus defined as a set of consecutive “jumps” {Δ*x*_*i*_} and “waiting times” between consecutive jumps {Δ*t*_*i*_}.

To simplify, we assume that *p*(Δ*x*) follows a Gamma distribution (Fig. 1B,C) of average *µ* and standard deviation 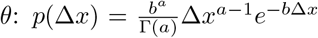with *µ* = *a/b* and 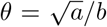. This distribution phenomenologically describes the typical distributions of loop extrusion steps observed experimentally with magnetic tweezers for condensin-mediated loop extrusion [31] (Fig. 1B). Interestingly, such a shape is characteristic of a multi-step exponential process [68] that may reflect the intrinsic stochastic kinetics of the capture of loop increments Δ*x* [36, 37].

### B. Investigated chromatin landscapes

In this work, we consider three types of accessibility describing the chromatin landscape faced by LEFs (Fig.1D): (1) homogeneous landscapes with uniform accessibility (0≤ *α*(*x*) ≡ *α* ≤1, ∀ *x*) ; (2) periodic landscapes characterized by a regular alternation of roadblocks of size *s* (e.g., nucleosomes) where the hinge cannot jump on (*α*(*x*) = 0) and of fully accessible regions of size 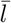(e.g., linker DNAs) with *α*(*x*) = 1 ; (3) random landscapes where roadblocks of size *s* are randomly positioned along the lattice (see below) and interspaced by fully accessible regions with a mean inter-obstacle distance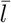.

In the latter case, random patterns are generated by emulating the possible random positions of an ensemble of non-overlapping particles on a unidimensional array, known also as a Tonk gas [69], which is a good model for nucleosome positioning [70]. This is performed by (1) randomly and uniformly partitioning the total number of fully accessible sites 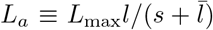 into *n* + 1 segments {*A*_*i*_}(∑ _*i*_ *A*_*i*_ = *L*_*a*_) with 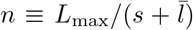 the corresponding number of roadblocks; and (2) inserting one roadblock between each segment. Note that we authorize *A*_*i*_ = 0, ie to juxtapose roadblocks with no accessible interspace. In the limit of large *L*_*a*_ and *n*, the distribution of interspaces *l* follows a geometric-like distribution of average 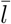 (Fig.1E, top) and the distribution of the number *m* of consecutive roadblocks a zero-truncated Poisson distribution of parameter 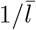 (Fig.1E, bottom) (See Supplementary Notes, Sec.I).

In Sections III A 4 and III B 1, in random landscapes, we allow the binding of the hinge to a site only if it belongs to a segment of consecutive fully accessible sites containing at least *bp*_min_ sites. This is equivalent to force *α*(*x*) = 0 for regions not satisfying such a criterion, effectively merging nearby roadblocks into larger composite barriers.

In Section III B 1, in periodic landscapes, before each simulated trajectory, we allow each roadblock to be stochastically removed with a probability *α*_*d*_. If removed, *α*(*x*) is set to 1 at the sites encompassing the ejected roadblock.

### C. Simulations and observables

For a fixed set of parameters, we simulate 10,000 trajectories, each during a fixed runtime *t*_max_ = 100*/k*_0_ and along a lattice of size *L*_max_ = 70, 000. Initial positions *x*_0_ are drawn uniformly within an accessible region around *x* = 10,000 to minimize boundary effects. In the case of random landscape, each trajectory is run on a different stochastic realization of the roadblock pattern (see above).

From the trajectories, we compute the average displacement⟨*x* −*x*_0_⟩ (*t*) from which we estimate the average speed *v*_mean_ from an affine fit (⟨*x* −*x*_0_⟩ (*t*) ≡*v*_mean_*t* +*b*_0_) over the range *t* = [0 : *t*_max_]. Similarly, we determine the initial velocity *v*_init_ from a linear fit but only over the time window *t* = [0, 5*/k*_0_]. The late-time scaling *w*_*f*_ is defined by fitting ⟨*x* −*x*_0_⟩ (*t*) by 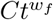 for *t* = [0.8*t*_max_, *t*_max_], performed via a linear fit in log-log scale. For example, in the case where ⟨*x* −*x*_0_⟩ (*t*) is a linear function of *t* over the full investigated time range, *v*_mean_ = *v*_init_ = *C, b*_0_ = 0 and *w*_*f*_ = 1. From the set of trajectories, we also compute the distribution of “instantaneous” speeds defined as the distribution of Δ*x*_*i*_*/*Δ*t*_*i*_ observed along trajectories (e.g.,Fig.2C).

**FIG. 2.**
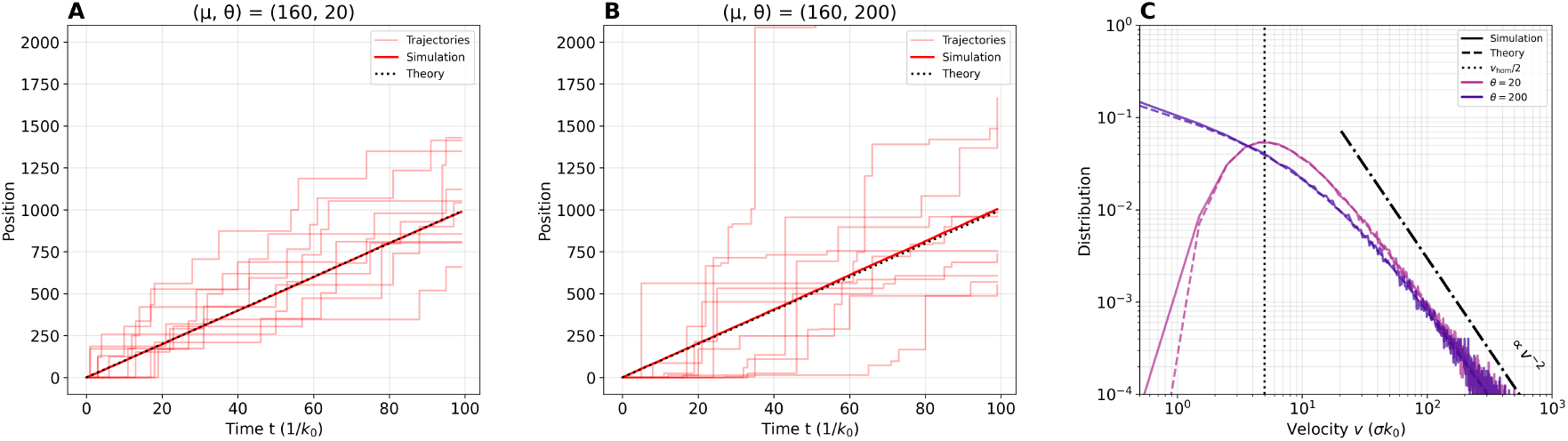
Loop extrusion kinetics along homogeneous landscapes for *α* = 10*/*160. **(A**,**B)** Examples of stochastic trajectories generated via the Gillespie algorithm (light red lines), and associated mean trajectory (thick red line) on *homogeneous* landscapes, for different values of (*µ, θ*), for *α* = 10*/*160 (*s* = 150, 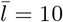 ). Simulation fits analytical results (dashed line).**(C)** Distributions *P* (*v*) of instantaneous velocities for the two parameter sets in (A,B) computed from simulations (full lines) and from analytical results (dashed lines). The vertical dotted line corresponds to 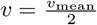, the dotted and dashed line to the scaling *P* (*v*) *∝ v*^*−*2^.

Note that, unless specified, times are given in 1*/k*_0_ unit, positions in *σ* unit and speeds in *σk*_0_ unit. Simulation code is available at https://github.com/npelletbossot/nucleo.

### D. Analysis of in vivo data

Positions of cohesin associated regions (CARs) of the budding yeast *Saccharomyces cerevisae* are inferred as in [71] by smoothing the raw signal with a Gaussian filter of width 500 bp and identifying the local peaks (peak value *>*10) along the genome of the Mcd1 cohesin subunit from ChIP-seq data taken from [72].

3C-like datasets are taken from recent PCP experiments in yeast, an alternative Micro-C-like method [73]. Raw contact matrices at 1 kbp resolution are treated using the <monospace>cooltools</monospace> library [74] as in [71]: first, they are ICE-balanced and then further distance-normalized by dividing them by the average contact frequencies *P* (*g*) for a given genomic distance *g*, leading to so-called observed-over-expected matrices representing the enrichment of contacts between any pair of loci *i, j* compared to the average value *P* (|*j* − *i*|).

Nucleosome properties in yeast are obtained by analyzing the nucleosome profile data from PCP experiments [73] using the DANPOS toolkit [75]. It infers nucleosome positions (peaks of the nucleosome signal) and gives for each nucleosome an intensity score (∼ height of the peak) that informs about the nucleosome occupancy and a fuzziness score (∼ width of the peak) that informs about the deviations of nucleosome position. Nucleosome repeat lengths (NRLs) are defined as the genomic distances between the centers (dyads) of nearest neighbor nucleosomes.

## III. Results

We develop a stochastic framework to model the growth of chromatin loop extruded by a one-sided loop extruding factor (LEF) like condensin (see Materials and Methods). In particular, we monitor the progression of the LEF hinge along chromatin by assuming that it occurs by consecutive stochastic jumps Δ*x* sampled from a Gamma-like distribution *p*(Δ*x*) at a given rate *k*_0_, as observed in vitro [31], and is hindered by the presence of roadblocks [40](Fig.1A). These roadblocks, on the other hand, can affect local chromatin compaction and modify the actual base-pair length captured by a single jump [37, 40].

### A. Chromatin accessibility plays major role in regulating loop extrusion speed

We start by investigating only the role of accessibility and roadblocks on loop extrusion dynamics and assume a constant homogeneous compaction (*c* = 1 *bp, σ* = 0.3*nm*) along the 1D array that models chromatin. At a given position *x*, accessibility is defined by *α*(*x*), which represents the probability that DNA at *x* is locally accessible to the LEF hinge for binding. For example, in the presence of a roadblock (like a nucleosome) at *x, α*(*x*) = 0 and an attempt to jump on *x* will be rejected.

#### 1. Homogeneous landscapes as a reference kinetic baseline

As a reference, we first consider the situation of constant homogeneous accessibility along chromatin (*α*(*x*) ≡*α*). This may represent a mean-field-like approximation where the roadblock random binding/unbinding dynamics to chromatin is very fast compared to loop extrusion kinetics; thus the chromatin fiber has an average accessibility 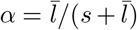 where *s* is the typical roadblock footprint on chromatin and 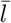 the average chromatin length between two consecutive roadblocks. In that simple case, analytical calculations can be made (see Supplementary Notes, Sec.II.A). The average position of the hinge scales linearly with time (⟨*x*(*t*) −*x*(*t* = 0) ⟩ = *k*_0_*αµt* with *µ* the average jump size) leading to an average extruding speed *v*_mean_ = *k*_0_*αµ*, independent of the fluctuations in jump sizes (characterized by *θ* the standard deviation of *p*(Δ*x*)) (Fig.2A,B). The distribution of instantaneous speeds is given by *P* (*v* ≡ Δ*x/*Δ*t*) = (*αk*_0_)*ab*(*vb*)^*a−*1^*/*(*αk*_0_ + *bv*)^*a*+1^ with *a* = (*µ/θ*)^2^ and *b* = *µ/θ*^2^ the parameters describing *p*(Δ*x*) (Fig.2C). For peaked jump distributions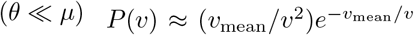is mostly driven by the distribution of waiting times Δ*t* between consecutive jumps and only depends on *v*_mean_ with an optimal value for *v* = *v*_mean_*/*2. For large jump distributions (*θ*≥ *µ*), *P* (*v*) is now dominated by the shape of the jump distribution, and the most probable instantaneous speed is 0.

#### 2 Periodic landscapes lead to phase-dependent kinetics

We now introduce the presence of fixed roadblocks, like nucleosomes, that prevent the binding of the hinge. In this section, we assume a periodic pattern of roadblocks (*α*(*x*) = 0) of size *s* = 150*σ*, interspaced by fully accessible linker sites (*α*(*x*) = 1) of size 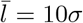 (Fig.1D center, see Materials and Methods). This type of regular spacing between obstacles may be representative of nucleosome positioning in constitutive heterochromatin regions or around the promoter of active genes [44, 46, 76]. For this configuration of roadblocks, we systematically vary the parameters of the jump-size distribution *p*(Δ*x*) to scan different situations with various means *µ* and standard deviations *θ*. All the simulated situations lead to linear average trajectory (Fig.S6) from which the average speed *v*_mean_ can be extracted.

For well-peaked jumps (*θ* ≪ *µ*), we observe a strong oscillatory dependence of *v*_mean_ to *µ* (Fig.3A, purple and blue lines, Fig.S6). When *µ* is about a multiple of 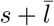 the period of the chromatin landscape, jumps are “in-phase” with accessible linkers, maximizing acceptance probability. In that case, *v*_mean_ approaches the maximum possible average speed *µk*_0_ and *P* (*v*) has the same shape as in the homogeneous case (see above) with a most probable speed *v* = *v*_mean_*/*2 (Fig.3C, purple line). In contrast, when 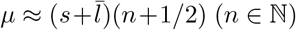, jumps are in “anti-phase” and repeatedly land on obstacles, trajectories stall, yielding *v*_mean_ ≈0 and the distribution of instantaneous speeds peaks at 0 (Fig.3C, pink line). Increasing the jump size dispersion (larger *θ* values) mitigates the “phase-locking” effect (Fig.3A, green and yellow lines) adding enough variability to allow hinge to escape the traps even if *µ* is in anti-phase. This comes at the cost of reduced maximum speeds, as oversized or undersized steps fail to optimally exploit the accessible linker regions. Therefore, when *µ* is “in-phase”, adding more dispersion in the jump size limits the average speed and *v*_mean_ is a decreasing function of *θ* (Fig.3B, purple line). In contrast, when *µ* is in “anti-phase”, more dispersion facilitates the hinge progression and *v*_mean_ is an increasing function of *θ* (Fig.3B, green and yellow lines). At large *θ* values, *v*_mean_ converges to *v*_hom_ ≡ *k*_0_ ⟨*α*(*x*) ⟩_*x*_*µ* where 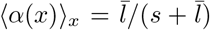 is the average accessibility along the 1D array (Fig.3B), as if the hinge were progressing in a partly-accessible homogeneous landscape. To fully compare with the corresponding homogeneous case, we plot the ratio *r*_*v*_ ≡ *v*_mean_(*µ, θ*)*/v*_hom_(*µ*) as a function of (*µ, θ*) (Fig.3D). This plot recapitulates the different oscillatory regimes and the presence of “in-phase” – fast (*r*_*v*_ ≫ 1) – and “anti-phase” – slow (*r*_*v*_ ≪ 1) – modes.

**FIG. 3.**
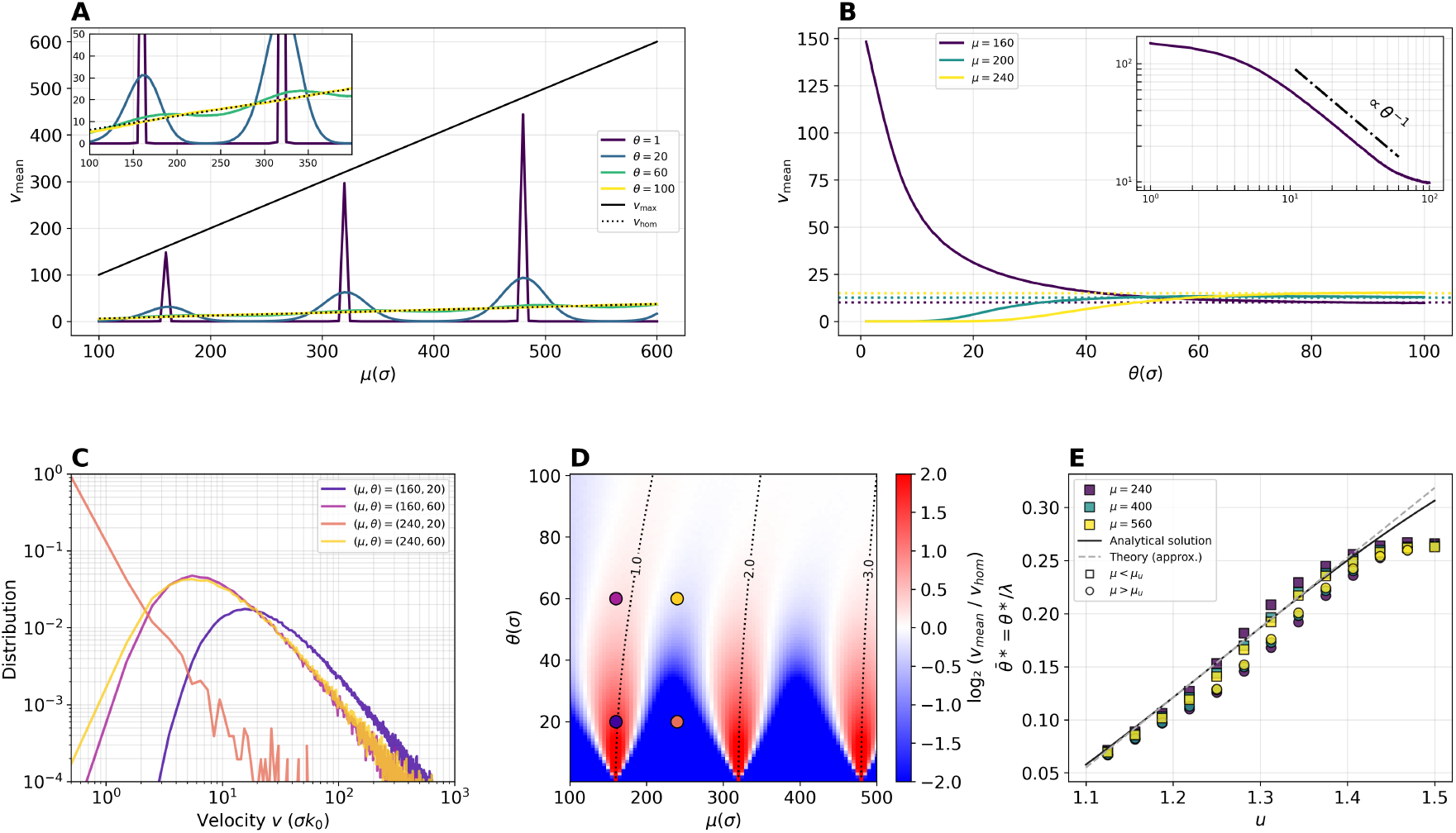
Loop extrusion kinetics along periodic landscapes for (*s* = 150, 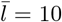 ). **(A, B)** Evolution of *v*_mean_ for a periodic landscape (*s* = 150, 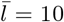 ) as a function of *µ* (A) and *θ* (B) for fixed values of *θ* and *µ*, respectively. Dotted lines represents the extrusion speed 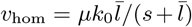 on corresponding homogeneous landscapes with same average accessibilities. The black line is the maximal reachable speed *v*_max_ = *µk*_0_ in absence of roadblock. Inset in (B) shows the curve for *µ* = 160 in log-log scale. **(C)** Distributions *P* (*v*) of instantaneous velocities for different values of (*µ, θ*) on a periodic landscape (*s* = 150, 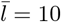 ). **(D)** Extrusion efficiency defined as the log2-ratio between *v*_mean_ and the corresponding *v*_hom_ as a function of *µ* and *θ*. The four points are the parameters used in panel (C). Dotted lines correspond to parameter values when the most probable jump size *µ − θ*^2^*/µ* is a multiple of 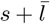 . **(E)** Normalized dispersion 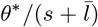 at which *v*_mean_ switches from low to high value as a function of the out-of-phase parameter *u*, where *u* = 1.5 indicates *µ* completely out-of-phase with the obstacle, whilst *u* = 1 is its counterpart, i.e., the in-phase scenario. Results are rescaled for different out-of-phase velocities.

A heuristic, semi-analytical theory can be derived for *v*_mean_ as ≈ *k*_0_ ⟨Δ*x*⟩ with ⟨Δ*x*⟩ the average size of accepted jumps (see Supplementary Notes, Sec.IIB and Fig.S1,S2). The theory is capturing very well the simulated results for over a large range of parameter values (Fig.S3). Under a low-noise approximation, we obtain analytically closed expressions for the variation of the velocity with the parameters. First, we analyze the optimal *µ* that locally maximizes *v*_mean_ for a fixed jump dispersion *θ*, finding that this is well described by a function of the form 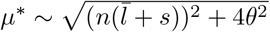 with 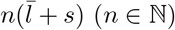 (*n* ∈ N) the periodic intervals between free linker regions (Fig.S4). Interestingly, the same dependency is also obtained when the most probable jump size *µ* − *θ*^2^*/µ* (i.e. the mode of *p*(Δ*x*)) is ““in-phase”, i.e. also a multiple of 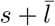 (dotted lines in Fig.3D). Next, we investigate the dependence of *v*_mean_ on *θ* for a fixed *µ*-value. In the in-phase case 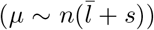, the velocity scales as *v*_mean_ ∼*θ*^*−*1^ over a wide range of *θ*-values (inset in Fig.3B and Fig.S5A). Between the in- and the out-of-phase case, i.e., 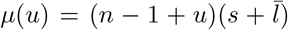 with *u*∈ [1, 1.5] (*u* = 1: in-phase, *u* = 1.5: anti-phase), we determined the characteristic dispersion level *θ*^*\**^ at which the system transitions from very low to high extrusion velocity *v*_mean_, typically the frontier between the blue and white/red regions in Fig.3D. We found that *θ*^*\**^ is proportional to 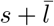 and can be described by a quasi-linear function of *u* (Fig.3E, and Fig.S5B).

#### 3. Random landscapes lead to non-linear loop extrusion dynamics

In this section, we consider irregular roadblock arrays where non-overlapping obstacles of size *s* = 150*σ* are randomly positioned along the array with fully-accessible linker inter-spaces of average size 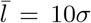 (Fig.1D,right, see Materials and Methods) to reach the same average accessibility than in the previous sections. However, in this scenario, roadblocks may be directly juxtaposed with no interspace, potentially leading to the rare formation of large effective obstacles (Fig.1E). This type of random spacing is characteristic of nucleosome positioning found in euchromatin, for example within gene bodies [46, 77].

Contrary to periodic patterns, the average trajectory of the hinge along the array may exhibit two regimes (Fig.4A, Fig.S7): a linear dynamics at short times characterized by an average speed *v*_init_ and a sub-linear power-law-like dynamics at longer time 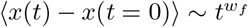 characterized by a kinetic exponent *w*_*f*_ *<* 1. This translates the behavior of individual stochastic trajectories (Fig.4A) showing the motion of the hinge until it stops for a very long time period in front of a large obstacle for which the probability of jumping over is very low. Although such large barriers are rare, the probability of encountering one increases as the hinge progresses along the array. Such an average sub-linear kinetics is characteristic of stochastic renewal processes with fat-tail distributions of waiting times between consecutive, productive events [78–80] (see Supplementary Notes, Sec.IIC). In our model, the presence of large obstacles (Fig.1E) coupled to *p*(Δ*x*) can lead to such a phenomenon (Fig.S8) and the distribution of instantaneous speeds can be bimodal (Fig.4C, pink line) with a strong peak for *v* = 0 and a weaker peak for *v* ≲ (*v*_mean_*/*2). As *θ* increases, the probability of jumping over large obstacles increases and *w*_*f*_ converges slowly to 1 (Fig.4B and Fig.S9).

**FIG. 4.**
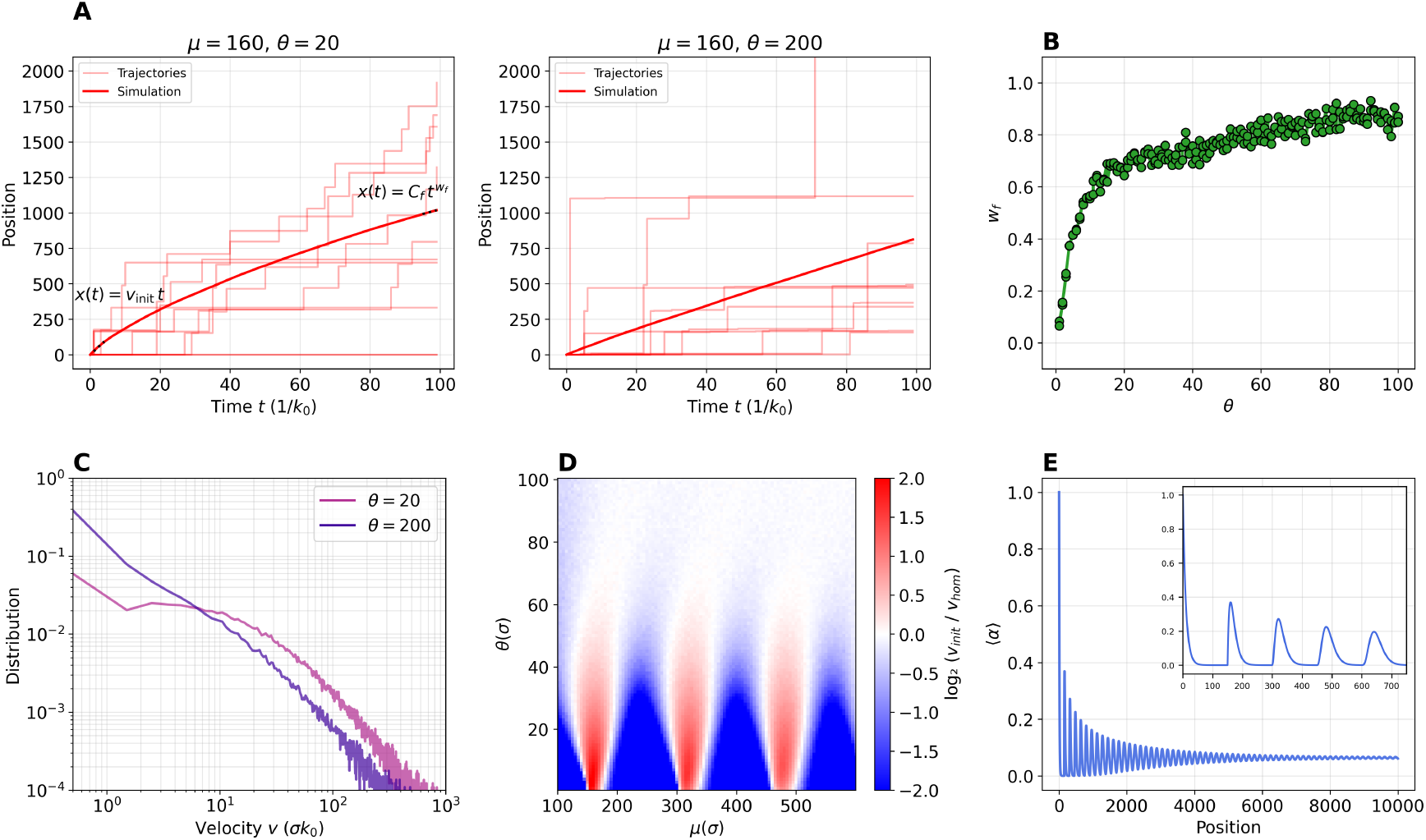
Loop extrusion kinetics along random landscapes for 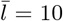. **(A)** Example of stochastic trajectories generated via the Gillespie algorithm (light red lines), and associated mean trajectory (thick red line) on *random* landscapes, for different values of (*µ, θ*). **(B)** Scaling law parameter *w*_*f*_ as a function of the jump size dispersion *θ* for *µ* = 160. **(C)** Distributions *P* (*v*) of instantaneous velocities for the two parameter sets of panel (A). **(D)** Extrusion efficiency defined as the log2-ratio between *v*_init_ and the corresponding *v*_hom_ as a function of *µ* and *θ*. **(E)** Mean accessibility profile *⟨α⟩* after a roadblock in *random* landscapes.

Interestingly, while roadblocks are *a priori* randomly positioned, *v*_init_ exhibits a qualitatively similar dependency on (*µ, θ*) than in the periodic scenario with in- and anti-phase behaviors (Fig.4D). This reflects the actual average accessibility “seen” by the hinge downstream an arbitrary roadblock (Fig.4E), which is characterized by dumped oscillations of period 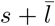 that persist over a wide range of sites. Indeed, these density oscillations result from statistical ordering and are classically observed for dense, one-dimensional hard-rod fluid [69] like nucleosomes along the chromatin [70]. However, in the in-phase regime, *v*_init_ is always smaller than in the corresponding periodic scenario with similar accessibility due to the presence of larger obstacles in the random scenario (Fig.S10). In contrast, in the anti-phase regime, random patterns lead to faster progression than periodic landscapes (Fig.S10).

#### 4. Roadblock density impacts hinge progression

In the previous sections, we characterized hinge progression for different roadblock landscapes, in the limit of dense arrays of roadblocks 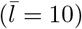 where only 6.25% of sites were accessible.

In the *periodic* landscape scenario, increasing the linker length 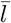 does not modify the phenomenology observed previously (Fig.5A and Fig.S6). Namely, for *θ* ≪ *µ*, we observe in-phase and anti-phase regimes when 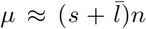 and 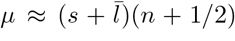 respectively. Increasing *θ* dumps the oscillatory behavior. Interestingly, when 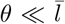, *µ*, the inherent asymmetric shape of the Gamma distribution *p*(Δ*x*) propagates into the shape of the in-phase regime (see the full gray line in Fig.5A).

**FIG. 5.**
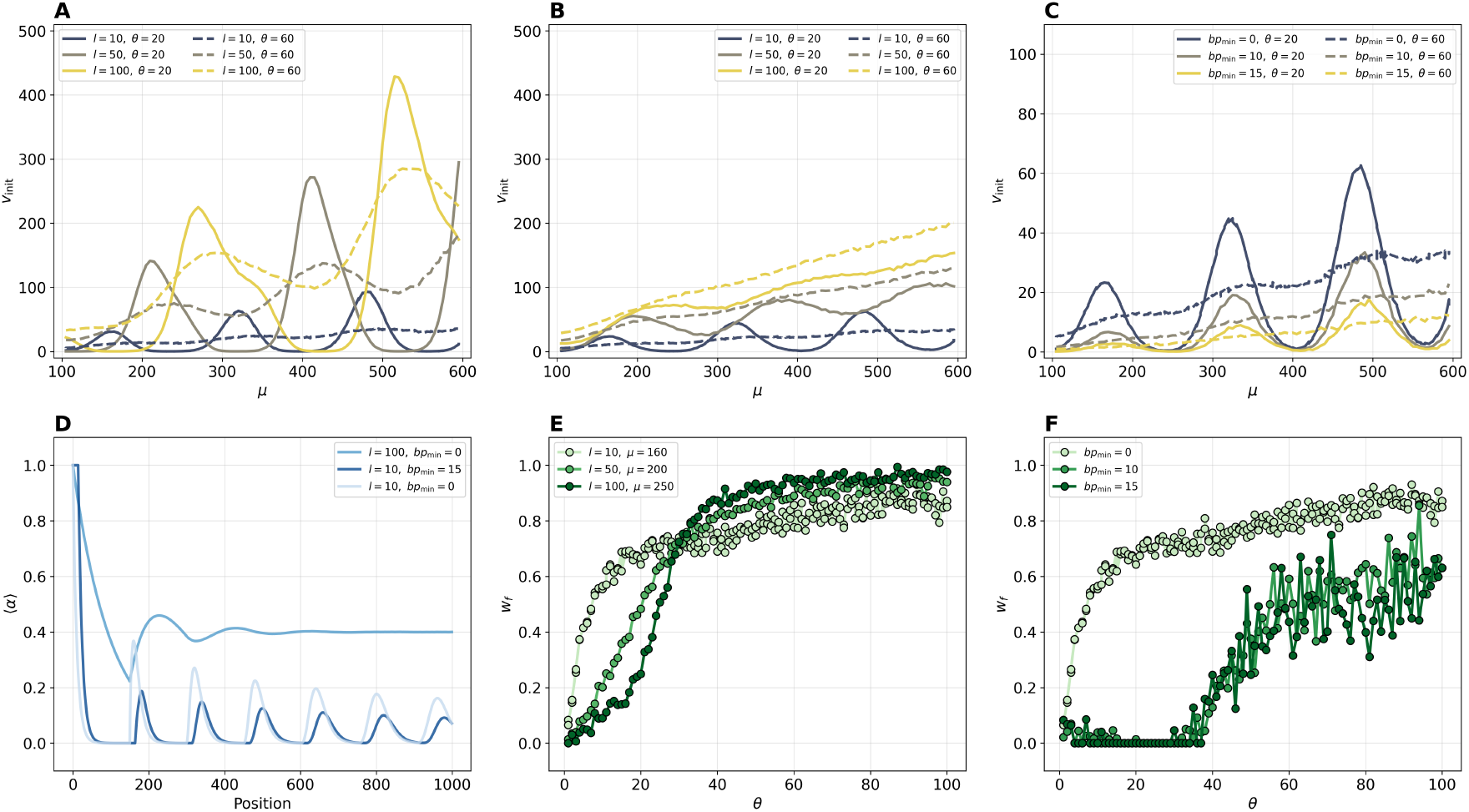
Impact of roadblock density (A,B,D,E) and minimal linker length (C,F) **(A)** Initial velocity *v*_init_ as a function of *µ* for different 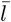 and *θ* values on *periodic* landscapes. **(B)** Initial velocity *v*_init_ as a function of *µ* for different 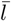 and *θ* values on *random* landscapes. **(C)** Initial velocity *v*_init_ as a function of *µ* for different *bp*_min_ and *θ* values on *random* landscape for 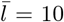. **(D)** Mean accessibility profile *⟨α⟩* after a roadblock in *random* landscapes for different combinations of 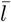 and *bp*_min_. **(E)** Scaling law parameter *w*_*f*_ as a function of the jump size dispersion *θ* for different (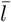; *μ*) combinations on *random* chromatin landscapes. **(F)** Scaling law parameter *w*_*f*_ as a function of the jump size dispersion *θ* for *µ* = 160 and different values of *bp*_min_ for 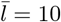.

Similarly, in the *random* landscape scenario, increasing the mean linker length 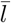 qualitatively maintains the same phenomenology as previously observed, but the reduction of roadblock clustering overall increases the jump acceptance rate and thus the hinge average speed *v*_init_ (Fig.5B and Fig.S9). The quasi-periodic behavior observed for *v*_init_ is present at low 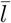 values, with the positions and widths of the in-phase and anti-phase regimes shifting as 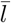 changes. However, these oscillations gradually dampen as 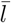 increases, a direct consequence of the gradual loss of roadblock statistical positioning (Fig.5D and Fig.S9). The system exhibits near-linear average trajectories (with *w*_*f*_ → 1) across a wider range of (*µ, θ*) values (Fig.S9) and converges more rapidly to a homogeneous-like behavior when *θ* increases (Fig.5E). However, in the in-phase regime, for low *θ* values, while *v*_init_ increases with 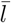 (Fig.5B), the parameter *w*_*f*_ decreases with 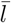 (Fig.5E). Indeed, for large 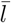, roadblocks are much less regularly positioned relative to one another (Fig.5D). Consequently, when the jump dispersion *θ* is not large enough to allow the hinge to bypass individual roadblocks, the probability of being blocked for a very long time by a particularly poorly placed obstacle—even in the “in-phase” regime—is paradoxically increased.

Previous in vitro studies [40–42] suggested that the condensin hinge may not be able to bind to too small linker inter-spaces between obstacles. We explore the implications of a minimal footprint size for the hinge by limiting the jumps to accessible segments containing at least *bp*_min_ consecutive free sites (Fig.S11). This is equivalent to set *α*(*x*) = 0 for regions not satisfying such a criterion. Even modest increases in *bp*_min_ prune short accessible gaps and merge adjacent roadblocks into larger effective composite barriers (Fig.S12). As a consequence, the probability to encounter very large obstacles is increased, leading to lower initial speed (Fig.5C) and reinforced sub-linear - confined - mean trajectories (Fig.5F).

### B. The interplay between compaction and roadblock density lead to optimal loop extrusion speed on in vivo chromatin

To focus on the role of accessibility, in the first part of this work, we assumed that the presence of road-blocks did not affect the local compaction of chromatin and imposed a constant homogeneous compaction along the 1D array (*c* = 1 *bp* per site). However, in vivo, the compaction of chromatin fiber is highly heterogeneous with a low compaction for linker DNA regions (∼ 1*bp/*0.3*nm*) and a high compaction for nucleosomal DNA (∼150*bp/*10.5*nm*, assuming one nucleosome contains ∼150*bp* and measures ∼10.5*nm*). Therefore, while the presence of roadblocks such as nucleosomes certainly limits chromatin accessibility (see above) and thus possibly alters loop extrusion dynamics, it may also “accelerate” hinge progression as a jump of a given physical size (in nm) over one or several roadblocks may contain a larger number of base-pairs than over a bare DNA segment of equivalent physical length.

To investigate this interplay between compaction and accessibility, we consider composite landscapes made up of linker and nucleosomal DNA regions: each site *x* of the array is of size *σ* = 0.3*nm* and contains either *c*(*x*) ≡ *c*_*l*_ = 1*bp/site* if within a linker DNA or *c*(*x*) ≡*c*_*n*_ = 4.3*bp/site* if within nucleosomal DNA regions. Each individual roadblock represents a nucleosome and has a footprint of *s* = 10.5*/*0.3 = 35 sites.

To model the hinge progression on chromatin, we first remark that estimations of the persistence length of in vivo chromatin (∼40 −60*nm*) [81–85] are very similar than the one of dsDNA (∼50*nm*). Therefore, as the bending rigidity of the underlying polymer likely plays a dominant role in defining the capacity of the hinge to jump [55, 64], we assume that the jump probability distribution function *p*(Δ*x*) in the chromatin context described here is the same as the one observed experimentally in vitro on naked dsDNA [31]. As a reference distribution, we consider the distribution of step sizes obtained by Ryu et al. [31] at 0.4 pN whose statistics is good enough to allow a fit by a Gamma-like distribution with *µ* = 37*nm* = 150*σ* and *θ* = 25*nm* = 100*σ* (Fig.1B). We will also vary these parameters to make sure our results do not depend on this arbitrary choice.

#### 1. The base-pair extrusion speed may be faster on chromatin than on naked dsDNA

We revisit the dependency of extrusion speeds as a function of linker sizes 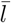 in the periodic and random landscape scenarios for (*µ, θ*) = (150, 100)*σ*. In particular, we study (1) the”site” loop extrusion speed 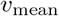, as defined previously and expressed in *σk*_0_ unit, and which does not account for the compaction ; (2) the “base-pair” loop extrusion speed 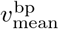 (in *bpk*_0_ unit) that integrates the local compaction and is given by 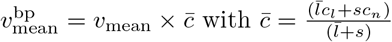 the average compaction along the array which is a decreasing function of 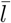 (Fig.6B).

**FIG. 6.**
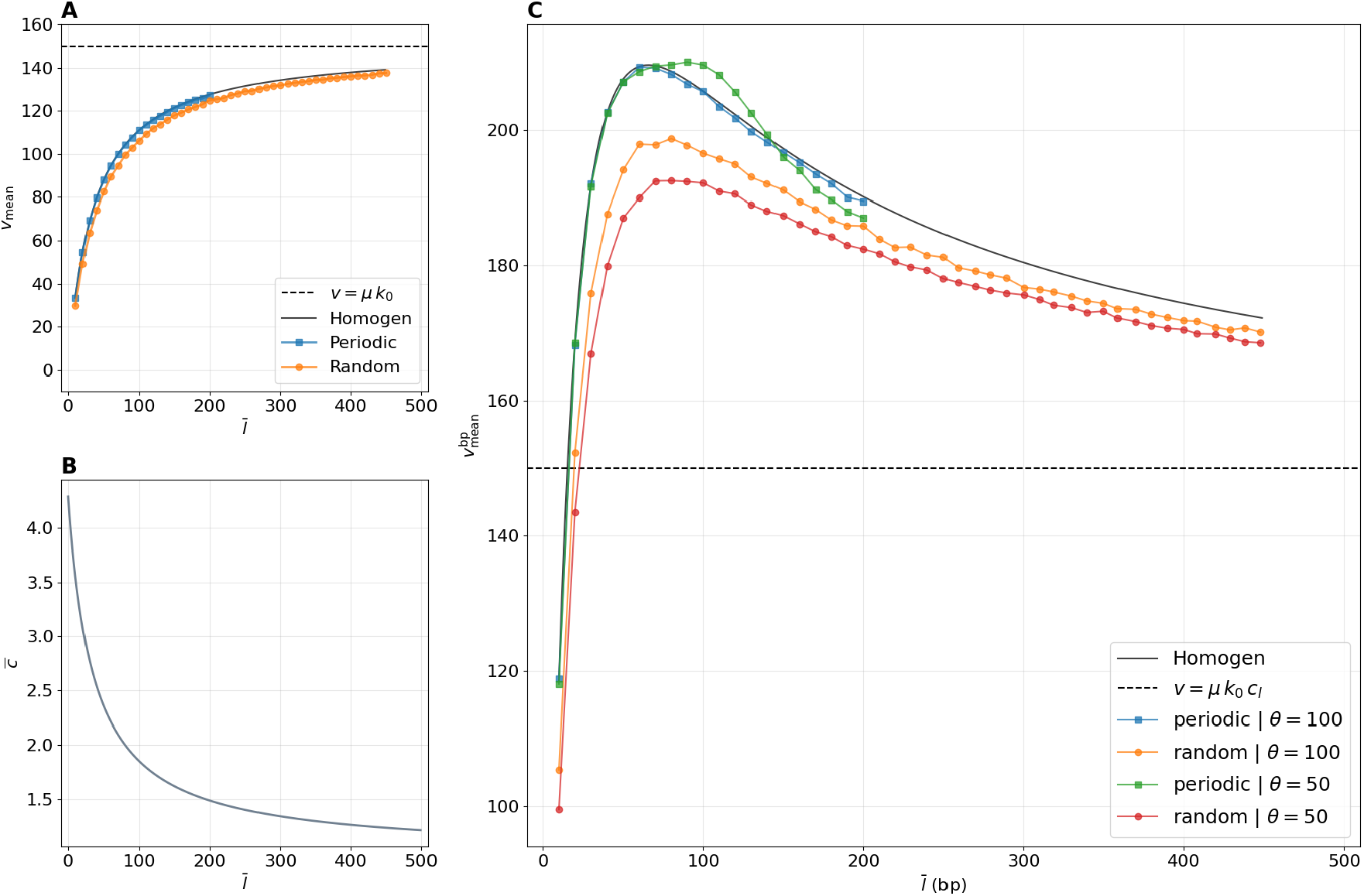
Impact of chromatin compaction on loop extrusion speed. **(A)** *v*_mean_ (in *σk*_0_ unit) as a function of 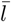 for (*µ* = 150, *θ* = 100) in the homogeneous, periodic and random scenarios. **(B)** Average compaction 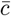 along the array. **(C)** 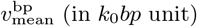 as a function of 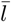 for *µ* = 150 and two different *θ* values in the homogeneous, periodic and random cases. The horizontal dashed line is the corresponding speed in the absence of roadblock.

We observe that *v*_mean_ (Fig.6A) is an increasing function of the linker length, due to rising accessibility as described in Sec.III A 4, and it converges to *µk*_0_, the speed on naked dsDNA for large linkers. Since the dispersion *θ* (= 100*σ*) of jump size is much greater than the roadblock size (*s* = 35*σ*), it is easier to bypass obstacles even for small linker lengths. Therefore, *v*_mean_ remains very close to *v*_hom_, the mean speed of the corresponding homogeneous case, in the periodic scenario and is slightly lower for random landscapes.

Strikingly, as 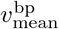 is the product between an increasing function of 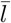 (*v*_mean_, Fig.6A) and a decreasing function of 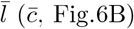 its evolution as a function of 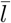 is non-monotonous (Fig.6C). In both periodic and random scenarios, the progression in *bp* along the chromatin became faster than on naked dsDNA (*µk*_0_) 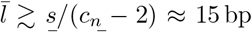 and is optimal for 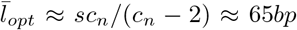 (see Supplementary Notes Sec.IIA). For 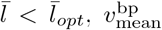 is mostly driven by chromatin accessibility, while, for 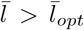, the compaction effect of nucleosomes dominates. Such a phenomenology is conserved for different values of *µ* and *θ* (Fig.6C and Fig.S13). In both types of landscape, increasing *θ* leads to an overall increase in 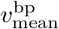, except in the periodic scenario with weak dispersion *θ* when 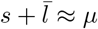, i.e. close to an in-phase regime (Fig.6C, green line).

When introducing a minimal binding site size *bp*_min_ in the random case (Fig.7A), 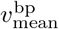 decreases significantly, in particular for low 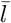, and the position of maximal velocity shifts to higher 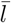 values. This translates the strong loss of accessibility that now dominates until larger 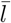 values.

**FIG. 7.**
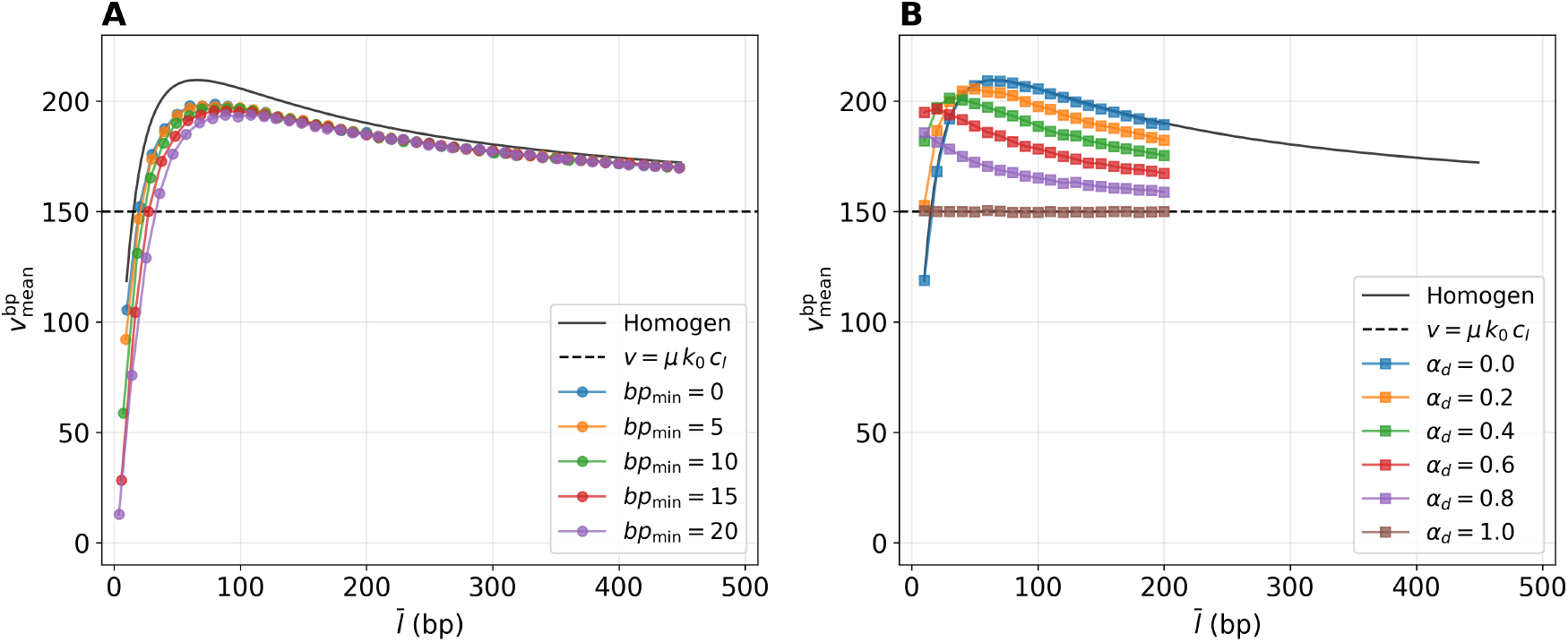
Impact of a minimal binding site size (A) and of nucleosome eviction (B). **(A)** 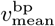 as a function of 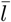 for (*µ* = 150, *θ* = 100) in the random case for various values of *bp*_min_. **(B)** 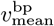 as a function of 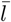 for (*µ* = 150, *θ* = 100) in the periodic case for various values of *α*_*d*_.

To simulate situations of well-positioned nucleosomes that are not always present but could have been removed by remodelers for example, we consider a periodic landscape with linker length 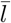 where we additionally randomly remove nucleosomes with a probability *α*_*d*_ prior to simulating each trajectory (Fig.7B). For *α*_*d*_ ≲ 0.5, at low 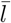, increasing *α*_*d*_ augments the bp hinge speed as accessibility improves. However, for higher 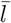 the reduction of compaction due to nucleosome removal dominates and limits the hinge progression in bp. As a consequence, the position of maximal velocity shifts to lower 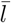 values. Interestingly, for *α*_*d*_ ≳ 0.5, compaction dominates in all occupancy regimes and 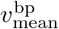 is a decreasing function of *α*_*d*_ for all 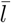 values.

#### 2. In vivo chromatin in yeast is dominated by compaction

Finally, we aim to challenge our model predictions with experimental data. We focus on budding yeast in the G2/M phase as loop extrusion by cohesin is well characterized at this cell cycle stage [71, 72, 86–88] and there exist several good quality datasets. In yeast, CARs (Cohesin-Associated Regions) are genomic sites enriched in cohesin and they are located between convergent genes [72]. On Micro-C, PCP or CAD-C contact maps, loops are observed between CARs [72, 73, 89] and disappear upon cohesin depletion, suggesting that these loops arise from loop extrusion and that CARs represent strong loop extrusion boundaries.

As there is no direct and systematic method to estimate loop extrusion speed in vivo so far, we develop an indirect strategy: if a LEF (cohesin) moves efficiently (in bp) along chromatin, it will cover the genomic distance between two neighboring CARs more quickly and thus will stabilize the loop between these CARs for longer and a stronger loop pattern should be visible on contact maps [61]. Our idea is therefore to correlate the contact enrichment between consecutive CARs with the properties of the chromatin fiber (nucleosome density, linker length, positional disorder) between them.

To do so, we infer the positions of CARs from cohesin ChIP-seq data [72], nucleosome properties (positions, intensity and fuzziness scores, nucleosome repeat lengths) from nucleosome data [89] and contact maps from PCP experiments [73] (see Materials and Methods). Then, for each pair of consecutive CARs *i, j*, we extract (Fig.S14) (1) the contact enrichment *I*_*o/e*_ as the average of the observed-over-expected (O/E) signal within a flanking window of ±1 kb around them, (2) the average cohesin signal *I*_*c*_ within a flanking window of ±1 kb around *i* and *j*, (3) the median nucleosome occupancy *I*_*α*_ defined as the median intensity score for the nucleosomes in between the CARs, (4) the median nucleosome repeat length 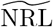 defined as the median spacing between consecutive nucleosomes in between *i* and *j* and (5) the median fuzziness 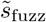 defined as the median fuzziness score for the nucleosomes in between *i* and *j* that quantifies the median positional disorder of individual nucleosomes in the region.

First, we verify that *I*_*o/e*_ is correlated with *I*_*c*_ independently of the genomic distance between consecutive CARs (Fig.8A, Fig.S15 A), supporting our working hypothesis: the more efficient the extrusion, the more quickly cohesins arrive at CARs, and therefore the greater the density of cohesin around CARs and stronger the enrichment in contact.

**FIG. 8.**
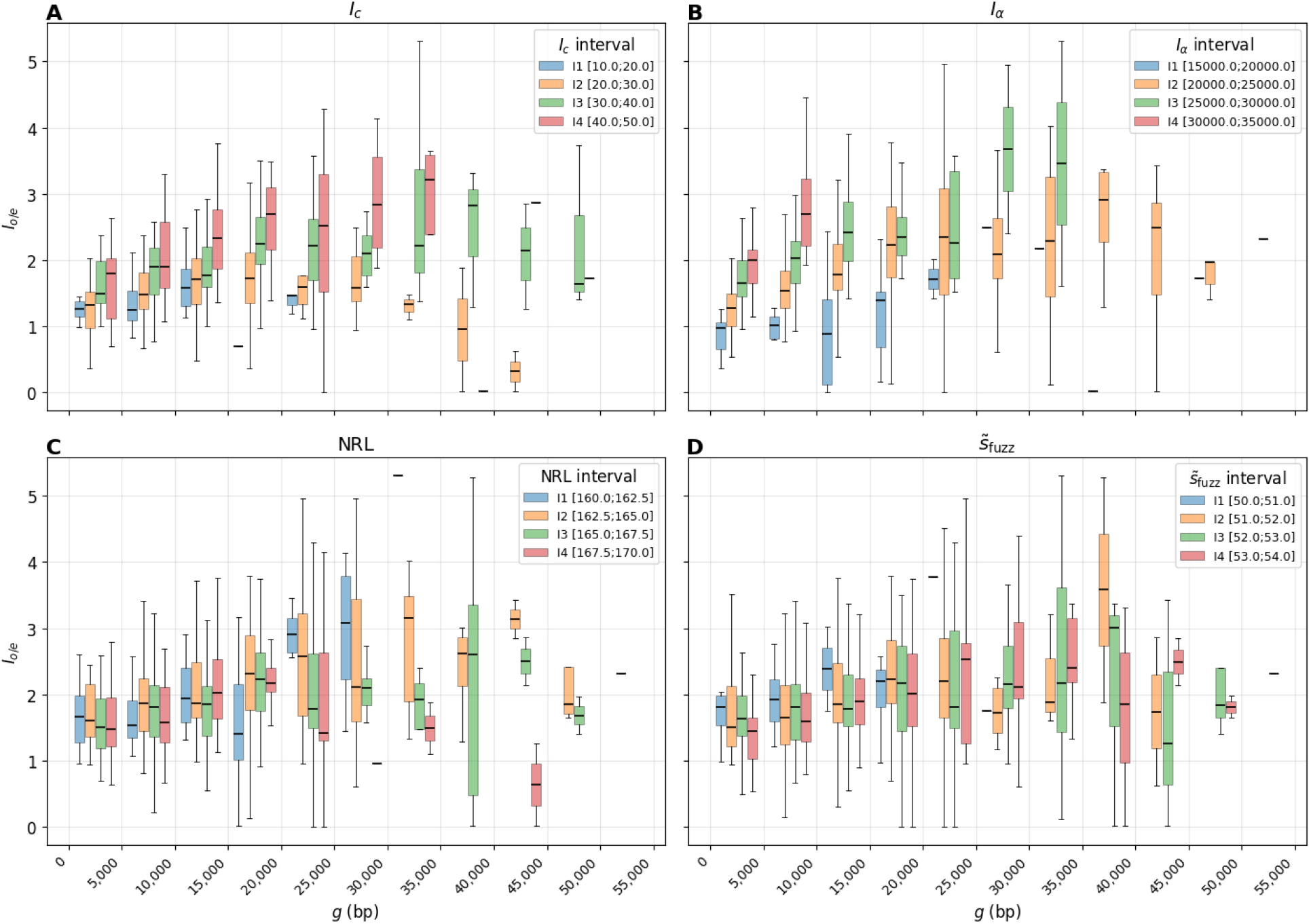
Correlation between the contact enrichment between consecutive CARs and the properties of the underlying nucleosomal landscape. **(A-D)** For each chromatin properties [*I*_*c*_ (A), *I*_*α*_ **(B)**, 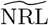 **(C)**, 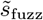 **(D)**], four equal-length intervals are defined covering the full range of values. All the pairs of consecutive CARs from a given interval are then grouped by genomic distances *g* between the CARs (5 kbp bins). Boxplots of *I*_*o/e*_ for each group are given for each property.

Then, we examine the relation between *I*_*o/e*_ and *I*_*α*_, a proxy for nucleosome occupancy in the region (Fig.8B, Fig.S15 B). For all the ranges of genomic distances between consecutive CARs, we observe a strong correlation between both quantities: the more nucleosome occupancy the more contact enrichment and thus putatively the higher 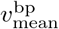.

We perform similar comparison with 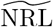 and find a weak anti-correlation only at large genomic distances (*>* 20 kbp) (Fig.8C, Fig.S15 C). It should be noted that the variability of 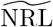 in yeast is relatively low (Fig.S14), with values ranging from 160 to 170 bp. Thus, if a minor effect of 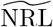 exists, it would be more readily detectable over large genomic distances, where such effects can accumulate.

Finally, we compare *I*_*o/e*_ and 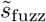 (Fig.8D, Fig.S15 D). The effects here appear very weak, and no clear trend seems to emerge, other than a possible—though non-significant—anti-correlation at short genomic distances. Here, too, the dispersion of fuzziness score is quite low (Fig.S14), yet no clear correlation is observed at long genomic distances.

Overall, contact enrichment is positively correlated with nucleosome occupancy, possibly anti-correlated with typical NRL, and no trend is observed for fuzziness. Translating into the language of our modeling, this implies that (1) decreasing *α*_*d*_ (=increasing nucleosomal occupancy via *I*_*α*_) is associated with greater bp extrusion speeds 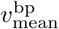, (2) that increasing 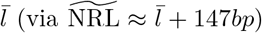 could be linked to a decrease in 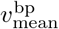; (3) that the dispersion of fuzziness (random vs. periodic landscape) is likely not large enough to observe an effect. In yeast, the fuzziness scores and inter-nucleosome distances show low dispersion, suggesting that the majority of the genome has a chromatin landscape close to a periodic or weakly random organization with a median linker 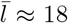 bp. Moreover, recent single-molecule measurements of nucleosome positioning in yeast [90] show that the average accessibility on chromatin is 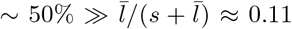. This implies that about half of the nucleosome positions are not occupied, equivalent to having *α*_*d*_ ≈ 0.5 in our modeling. In this regime, our model predicts that the main effect of nucleosome occupancy on 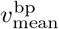 is driven by the compaction role of nucleosomes rather than by their impact on accessibility (Fig.7). Thus, increasing 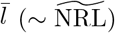 or 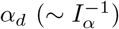 should reduce 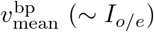 as observed in our analysis of experimental data.

## IV. Discussion and conclusion

In this work, we developed a minimal stochastic framework coupling a Gamma-distributed stepping process to an explicit description of the chromatin accessibility landscape.

The model reveals a rich dependence of extrusion dynamics on the linear pattern of accessibility and on the statistical properties of the jumps. Periodic landscapes may generate in-phase and anti-phase regimes, with efficient progression when the characteristic jump size is compatible with the roadblock spacing. Random landscapes, surprisingly, may preserve these patterns for short linker lengths, although they consistently exhibit lower extrusion efficiency. The presence of large obstacles or of a minimal engagement length (*bp*_min_) can significantly slows down extrusion, notably leading to the emergence of sub-linear regimes. These observations suggest that, for extrusion to proceed efficiently despite the heterogeneity of the chromatin landscape, the distribution of jump sizes (*θ* in our approach) must be sufficiently broad to allow for the exploration of a wide range of accessible positions. Together, these results show that the chromatin architecture acts as a filter on the intrinsic stepping dynamics of the motor rather than simply being a passive obstacle.

A central result of this work is that chromatin compaction can fundamentally change extrusion efficiency. While nucleosomes reduce local accessibility and therefore constrain the progression of the hinge in physical space, they also pack more base pairs into a given physical distance. Consequently, a jump of fixed size in nanometers can correspond to a larger genomic displacement on compacted nucleosomal DNA than on linker DNA. In both periodic and random landscapes, this leads to a competition between two opposing effects due to the presence of nucleosomes: increasing linker length improves accessibility and therefore facilitates hinge progression, while simultaneously reducing the average compaction and thus the number of base pairs covered by a jump of given physical size. This crosstalk is similar, even if not formulated the same way, to effects observed by Maji et al. [63] who modeled the diffusion of cohesin on nucleosomal landscape. In our case, the extrusion speed expressed in base pairs 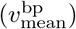 is thus a non-monotonic function of linker length and may reach an optimum at intermediate linker lengths (around 10 − 70 *bp*), depending on the minimal engagement length (*bp*_min_) and on the nucleosome eviction rate (*α*_*d*_). An interesting consequence is that the extrusion speeds measured *in vitro* on naked DNA and *in vivo* should be of the same order of magnitude.

This is consistent with recent in vivo estimations using live imaging 0.5 − 3 kbp/s [14, 91], equivalent to the speeds observed in vitro [25, 29].

Our analysis of yeast data demonstrates that nucleosome occupancy is significantly correlated with higher contact enrichment between extrusion boundaries, and thus with more efficient progression in base-pair. Even if we cannot fully rule out possible biological confounding factors like the role of transcription or technical biases from the experimental protocols, within our framework, this suggests that compaction effects may dominate in this organism. As in vivo data with endogenous chromatin landscape could be quite noisy (Fig.S15), future dedicated experiments, where the loading of LEFs [61] and the properties of the nucleosome positioning can be controlled, are needed to properly disentangle such a dual role of nucleosomes.

Although our minimal framework allows to describe how the mesoscopic extrusion dynamics emerge from the interplay between the “microscopic” stepping extrusion statistics and roadblock positioning, it relies on several simplifying assumptions to facilitate the interpretability of the model.

First, our model is restricted to the case of asymmetric extrusion by a single, isolated condensin. Consequently, in its current form, our model cannot account for the possible collective and cooperative effects that may arise from the simultaneous action of multiple motors on a single substrate—whether in terms of competition for the same accessible segments or, conversely, mutual facilitation in overcoming obstacles. Similarly, we consider only the case of unidirectional – condensin-like – extrusion here, whereas cohesin is expected to work bidirectionally. Nevertheless, this latter case can be viewed as a straightforward extension of our formalism, in which there is an alternation of stochastic moves in opposite directions according to the same rules [19].

Second, the accessibility landscape is assumed to be static: nucleosomes and roadblocks remain fixed throughout each simulation. As long as their intrinsic binding/unbinding or turnover dynamics (about hours for nucleosomes [92]) is slower than loop extrusion (second to minute scale), this approximation remains valid unless remodeling factors may actively impinge on LEF progression [63, 93] along chromatin as observed for the transcription machinery [94].

Finally, in our framework, chromatin is represented as a one-dimensional lattice on which the condensin hinge performs stochastic jumps. This description neglects the fine-scale three-dimensional organization of the chromatin fiber which may modify the mechanical environment experienced by the extruding complex. In particular, we considered jump distributions that were assumed constant and calibrated on in vitro experiments on naked DNA [31]. Even if we showed the robustness of our conclusions regarding the values of parameters (*µ, θ*), it is unclear how the presence of roadblocks may actually impact jump distribution by effectively changing the local chromatin fiber mechanical properties [40, 55, 64, 95].

Overall, the present framework establishes chromatin accessibility and compaction as strong, tunable determinants of loop-extrusion efficiency. Chromatin architecture can consequently be viewed not merely as a constraint on loop extrusion, but as a control parameter capable of regulating its dynamical regime. Moreover, our approach provides a modular basis to analyze experimental data as we illustrated, in yeast, showing the importance of chromatin compaction in loop extrusion.

## Acknowledgments

We thank Aurèle Piazza, Cédric Vaillant, Enrico Carlon, Je-Kyung Ryu, Stéphane Marcand and members of the Jost lab for fruitful discussions. We acknowledge Agence Nationale de la Recherche (Grant No.ANR-22-CE12-0035 fo P.B. and D.J.) for funding and Centre Blaise Pascal de simulation et modelisation numerique of the ENS de Lyon for computing resources. Work in the Jost lab is supported by government grants managed by the Agence Nationale de la Recherche under the France 2030 program, with the reference numbers ANR-24-EXCI-0001, ANR-24-EXCI-0002, ANR-24-EXCI-0003, ANR-24-EXCI-0004, and ANR-24-EXCI-0005.

