## Supplementary Notes and Figures for "Understanding the role of chromatin accessibility and compaction on loop extrusion dynamics"

### NOTATION

| Symbol | Parameter |
| --- | --- |
| $\bar{l}$ | Average linker size |
| $s$ | Nucleosome dimensions |
| $\lambda$ | Wavelength ($\bar{l} + s$) of the nucleosome-linker organization |
| $\mu$ | Mean jump length |
| $\theta$ | Standard deviation of the jumps |
| $a$ | Shape parameter of the Gamma distribution function |
| $b$ | Rate parameter of the Gamma distribution function |
| $k_0$ | Characteristic frequency of jump attempts |

### I. DISTRIBUTION OF INTERSPACES AND ROADBLOCKS IN THE RANDOM LANDSCAPE SCENARIO

In our simulations, we aim to generate random patterns of total length $L_{max}$ containing $n \equiv L_{max}/(s + \bar{l})$ roadblocks of size $s$ and $L_a \equiv L_{max}\bar{l}/(s + \bar{l})$ accessible sites, where the accessible sites are randomly distributed in between roadblocks (or upstream of the first roadblock or downstream of the last roadblock of the array), leading to $n + 1$ segments of size $\{A_i \geq 0, i \in [1 : n + 1]\}$ ($\sum_i A_i = L_a$). Note that we authorize $A_i = 0$, i.e., to juxtapose roadblocks resulting in no accessible interspace. Practically, this is done by (1) randomly and uniformly partitioning $L_a$ into $n + 1$ parts $\{A_i\}$; and (2) then build the full array by “inserting” one roadblock between each segment.

The probability to insert a given roadblock in between two sites is $p \equiv 1/(L_a + 1)$. Therefore the probability to have $m \geq 1$ consecutive roadblock inserted at the same location follows a binomial-like distribution $P(m) = C_n^m p^m (1 - p)^{n-m} / (1 - (1 - p)^n)$, where $C_n^m$ is the binomial coefficient. In the limit of large $L_a$ ($p \ll 1$) and $n$, $P(m)$ become equivalent to a zero-truncated Poisson distribution of parameter $np \approx 1/\bar{l}$, and for $m \geq 1$,

$$P(m) = \frac{1}{\bar{l}^m (e^{1/\bar{l}} - 1)m!} \quad (1)$$

The probability to have a linker inter-space of size $l = 0$ is given by $P(0) = \sum_{m=1}^{\infty} G(m)(m/(m+1)) \approx 1 - A$ where $G(m) = C_n^m p^m (1 - p)^{n-m}$ is the probability to find an obstacle containing $m$ consecutive roadblocks, $m/(m+1)$ is the number of junctions (i.e., of inter-space of size 0) between these roadblocks and $A = \bar{l}(1 - e^{-1/\bar{l}})$. Thus, the probability to find a linker of size $l \geq 1$ is given by $P(l) = AG(m=0)e^{l-1} \sum_{m \geq 1} G(m) \approx A(e^{1/\bar{l}} - 1)e^{-l/\bar{l}}$.

### II. ANALYTICAL THEORY OF HINGE PROGRESSION

#### A. Homogeneous landscapes

For a homogeneous landscape of accessibility $\alpha$, the probability $P(x, t)$ for the hinge to be at position $x$ at a given $t$ follows the master equation

$$\frac{dP(x, t)}{dt} = k_0\alpha \sum_{\Delta x=1}^{x-x_0} p(\Delta x)P(x - \Delta x, t) - k_0\alpha P(x, t) \quad (2)$$

with $x_0$ the initial hinge position at $t = 0$. From Eq. 2, we can compute the evolution equation of $\langle x \rangle(t) \equiv \sum_x xP(x, t)$:

$$\begin{aligned} \frac{1}{k_0\alpha} \frac{d\langle x \rangle(t)}{dt} &= \sum_{x=x_0}^{\infty} \sum_{\Delta x=1}^{x-x_0} xp(\Delta x)P(x - \Delta x, t) - \sum_{x=x_0}^{\infty} xP(x, t) \\ &= \sum_{v=1}^{\infty} \sum_{u=x_0}^{\infty} (u+v)p(v)P(u, t) - \langle x \rangle(t) \\ &= \langle \Delta x \rangle \equiv \mu \end{aligned} \quad (3)$$

leading to $\langle x - x_0 \rangle(t) = k_0\alpha\mu t = v_{\text{mean}}t$ with $v_{\text{mean}} = k_0\alpha\mu$.

Following the exact Gillespie algorithm we implemented, the distribution $p_w$ of waiting time $\Delta t$ between two consecutive jumps is exponentially distributed ($p_w(\Delta t) = \alpha k_0 \exp(-k_0\alpha\Delta t)$). Therefore, the distribution $P(v)$ of instantaneous speed $v = \Delta x/\Delta t$ is given by

$$\begin{aligned} P(v) &= \int_0^{\infty} d\Delta t \int_0^{\infty} d\Delta x p_w(\Delta t) p(\Delta x) \delta(v - \Delta x/\Delta t) \\ &= \int_0^{\infty} d\Delta t p_w(\Delta t) \Delta t p(v\Delta t) \\ &= \frac{\alpha k_0 b^a v^{a-1}}{\Gamma(a)} \int_0^{\infty} d\Delta t \exp[-(k_0\alpha + bv)\Delta t] (\Delta t)^a \\ &= \frac{\alpha k_0 ab(vb)^{a-1}}{(\alpha k_0 + vb)^{a+1}} \end{aligned} \quad (4)$$

with $\delta(x)$ the Dirac-delta function and $p(\Delta x) = \frac{b^a}{\Gamma(a)} \Delta x^{a-1} e^{-b\Delta x}$ the probability distribution of jumps (of average $\mu$ and standard deviation $\theta$) where $a = (\mu/\theta)^2$ and $b = \mu/\theta^2$. The mode of $P(v)$, i.e., the most probable positive $v$-value, is given by solving $\partial P/\partial v = 0$. From Eq. 4, this leads to

$$v_{\text{opt}} = \frac{(a-1)\alpha k_0}{2b} = \frac{(a-1)v_{\text{mean}}}{2a} \quad (5)$$

which is positive only if $a \geq 1$ (i.e., $\theta \leq \mu$). When, $\theta > \mu$, $v_{\text{opt}} = 0$.

In the limit of peaked $p(\Delta x)$, i.e., $\theta \ll \mu$ or $a \gg 1$, $p(\Delta x) \approx \delta(\Delta x - \mu)$ and $P(v) \approx \int d\Delta t p_w(\Delta t) \Delta t \delta(v\Delta t - \mu) = (v_{\text{mean}}/v^2) e^{-v_{\text{mean}}/v}$ and $v_{\text{opt}} = v_{\text{mean}}/2$.

By definition, $\alpha = \bar{l}/(s + \bar{l})$, therefore, $v_{\text{mean}}^{\text{bp}}$ is given by

$$v_{\text{mean}}^{\text{bp}} = v_{\text{mean}} \bar{c} = \mu k_0 \frac{\bar{l}c_l + sc_n}{\bar{l} + s} \frac{\bar{l}}{\bar{l} + s}. \quad (6)$$

Solving $v_{\text{mean}}^{\text{bp}} = \mu k_0$ gives $\bar{l} = s/(c_n - 2)$, the value above which $v_{\text{mean}}^{\text{bp}}$ is greater than extrusion velocity on naked DNA. Solving $\partial v_{\text{mean}}^{\text{bp}}/\partial \bar{l} = 0$ leads to $\bar{l}_{\text{opt}} = sc_n/(c_n - 2)$ the linker length where $v_{\text{mean}}^{\text{bp}}$ is optimal.

#### B. Heuristic model for periodic landscapes

We aim to determine the mean velocity of the hinge jumping on a periodic trail of roadblocks. In the long time limit, we expect that every position within the free space located between two obstacles is equally likely to be occupied. This is a reasonable hypothesis, which can be justified by considering the stochastic nature of the jumps.

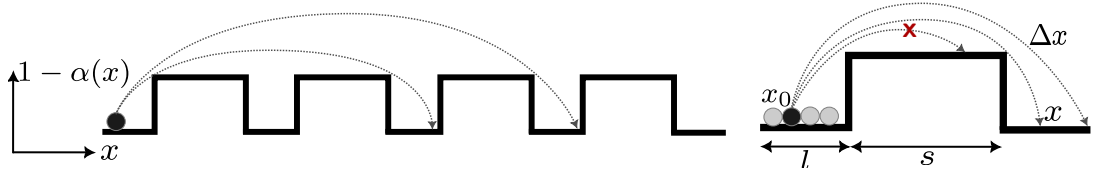

**Figure S1:** Heuristic model. A motor located at position $x_0$ can jump to position $x = x_0 + r$ with a probability given by the probability of making the jump and the probability of accepting the jump, which depends on the occupancy of the landing position, $\alpha(x)$.

A motor located at position $x_0$ makes a jump of length $\in [\Delta x, \Delta x + d\Delta x]$ with probability $p(\Delta x)d\Delta x$, see scheme in Figure S1). The jump is accepted or not depending on the occupancy of the landing position $x = x_0 + \Delta x$, given by $\alpha(x)$. For the initial position $x_0$, the average landing location $x$ is thus:

$$\begin{aligned}\langle x|x_0 \rangle &= x_0 + \langle \Delta x|x_0 \rangle \\ &= x_0 + \int_0^\infty \alpha(x_0 + \Delta x)p(\Delta x)\Delta x d\Delta x.\end{aligned}\quad (7)$$

Averaging over every possible initial position within the free space before the obstacle $x_0 \in [0, \bar{l}]$ (Figure S1),

$$\begin{aligned}\langle y \rangle &= \langle x|x_0 \rangle = \langle x_0 \rangle + \langle \Delta x|x_0 \rangle \\ &= \frac{1}{\bar{l}} \int_0^{\bar{l}} dx_0 \left[ x_0 + \int_0^\infty \alpha(x_0 + \Delta x)p(\Delta x)\Delta x d\Delta x \right] \\ &= \frac{1}{\bar{l}} \int_0^{\bar{l}} dx_0 \left[ x_0 + \int_{x_0}^\infty \alpha(x)p(x - x_0)(x - x_0)dx \right] \\ &= \frac{1}{\bar{l}} \int_0^{\bar{l}} dx_0 \left[ x_0 \left( 1 - \int_{x_0}^\infty \alpha(x)p(x - x_0)dx \right) + \int_{x_0}^\infty \alpha(x)p(x - x_0)x dx \right].\end{aligned}\quad (8)$$

This last equality has a clear probabilistic interpretation: the term $1 - \int \alpha(x)p(x - x_0)dx$ is the probability of not accepting the jump from $x_0$. Therefore, if the jump is not successful the particle remains at $x_0$, contributing to the mean position.

To determine the mean velocity $\langle v \rangle$, we simply remark that the final position $x$ found in the equation above is *per jump event*, thus $\langle v \rangle = k_0 \langle \Delta x \rangle k_0 \langle x - x_0 \rangle$, leading to

$$\langle v \rangle = \frac{k_0}{\bar{l}} \int_0^{\bar{l}} dx_0 \int_{x_0}^\infty \alpha(x)p(x - x_0)(x - x_0)dx. \quad (9)$$

To model the periodic obstacles we can use a trail of boxcar functions $\Pi$,

$$\begin{aligned}\alpha(x) &= 1 - \sum_{n=1}^\infty \Pi(x|s, l, n), \\ &= 1 + \sum_{n=1}^\infty \{ \mathcal{H}(x - n(\bar{l} + s)) - \mathcal{H}(x - n\bar{l} - (n - 1)s) \}\end{aligned}\quad (10)$$

where $\mathcal{H}$ is the Heaviside function. Hence,

$$\langle v \rangle = \frac{k_0}{\bar{l}} \int_0^{\bar{l}} dx_0 \left[ \int_{x_0}^{\bar{l}} p(x - x_0)(x - x_0)dx + \sum_{n=1}^\infty \int_{n(\bar{l} + s)}^{(n+1)\bar{l} + ns} p(x - x_0)(x - x_0)dx \right]. \quad (11)$$

The first integral in the parenthesis accounts for the movement before the first obstacle, and the term in the summation accounts for the free spaces between obstacles. We used Mathematica to solve the integral [1]. The resulting equation is lengthy but can be expressed as:

$$\langle v \rangle = k_0 \left( \mu - \frac{1}{b^2 \bar{l}} \frac{\Gamma(a + 2)}{\Gamma(a)} + \sum_i f_i \frac{\Gamma(a + 1, y_i)}{\Gamma(a)} + \sum_i g_i \frac{\Gamma(a + 2, y_i)}{\Gamma(a)} \right) \quad (12)$$

where

$$y_i \in \{ \bar{l}b, \quad n\lambda b, \quad (n\lambda - \bar{l})b, \quad (n\lambda + \bar{l})b \}, \quad (13)$$

the term $\lambda := \bar{l} + s$ is the periodicity of the landscape and $f_i$ and $g_i$ are functions of $n, \bar{l}, a, b$. The terms $\Gamma(\cdot, \cdot)$ are the lower incomplete Gamma functions.

#### B.1. Approximation in terms of elementary functions

The expression found (Eq. 12) captures the response observed in the simulations (Figure S3). However, the equation is too complex to provide a deep understanding of how the different parameters affect the velocity. Therefore, we search for approximations in terms of elementary functions.

First, we will omit for further calculations the result of the integral inside the bracket before the summation in Equation 11: since $\mu \gg \bar{l}$, we consider that the movement before the first obstacle is negligible. Next, we tackle the incomplete Gamma functions $\Gamma(v, x) \equiv \int_x^\infty dt t^{v-1} e^{-t}$. First, we re-express $\Gamma(a+1, y)$ and $\Gamma(a+2, y)$ following the relation $\Gamma(a+1, y) = x^a e^{-y} + a\Gamma(a, y)$. This results in terms of the form:

$$(n\lambda + \bar{l})^a, \quad (n\lambda - \bar{l})^a, \quad (14)$$

and in terms of the form $\Gamma(a, y)/\Gamma(a)$, which will be expressed using the regularized incomplete Gamma function $Q(a, y)$,

$$Q(a, y) \equiv \frac{\Gamma(a, y)}{\Gamma(a)}. \quad (15)$$

To simplify Gamma incomplete functions of the form $Q(a, (n\lambda b \pm \bar{l}b))$, we will Taylor expand around the small parameter $\bar{l}b \ll 1$,

$$Q(a, n\lambda b \pm \bar{l}b) \approx Q(a, n\lambda b) \pm Q'(a, n\lambda b) \bar{l}b \quad (16)$$

where

$$Q'(z, x) \equiv \partial_x Q(z, x) = -x^{z-1} e^{-x} \quad (17)$$

Hence,

$$Q(a, n\lambda b \pm \bar{l}b) \approx Q(a, n\lambda b) \mp \frac{(n\lambda)^{a-1} e^{-n\lambda b}}{\Gamma(a) b^{1-a}} \bar{l}b, \quad (18)$$

and, for the low noise scenario $\mu \gg \theta$, allow us to apply Stirling's approximation to the Gamma function for large $a$,

$$\Gamma(a) \approx a^a e^{-a} \sqrt{\frac{2\pi}{a}} \quad (19)$$

obtaining

$$Q(a, n\lambda b \pm \bar{l}b) \approx Q(a, n\lambda b) \mp \frac{(n\lambda)^{a-1} b^{1/b} e^{-b}}{b^{1-b} \sqrt{2\pi b^{-1}}} e^{-n\lambda b} \bar{l}b \quad (20)$$

which can be written in terms of the moments as

$$Q\left(\frac{\mu^2}{\theta^2}, (n\lambda \pm \bar{l}) \frac{\theta^2}{\mu}\right) \approx Q\left(\frac{\mu^2}{\theta^2}, n\lambda \frac{\theta^2}{\mu}\right) \mp \frac{\mu}{\sqrt{2\pi\theta}} \frac{\bar{l}}{n\lambda} \left(\frac{n\lambda e}{\mu}\right)^{\mu^2/\theta^2} e^{-\frac{\mu n\lambda}{\theta^2}}. \quad (21)$$

At this stage, we look to simplify the lower incomplete Gamma function $Q(z, x)$. For $z \sim x$, the incomplete Gamma function can be approximated by the Temme approximation [2, 3]:

$$Q(z, x) \approx \frac{1}{2} \operatorname{erfc}\left(\frac{x-z}{\sqrt{2z}}\right) \quad (22)$$

The complementary error function $\text{erfc}$ can be described phenomenologically by a logistic function of the form,

$$\text{erfc}(x) \approx \frac{2}{1 + \exp(Cx)}, \quad (23)$$

where $C$ is a constant. Assisted by ChatGPT, we constrained the phenomenological approximation to match the slope of $\text{erfc}$ at $x = 0$.

$$\left. \frac{d \text{erfc}(x)}{dx} \right|_{x=0} \stackrel{!}{=} \left. \frac{d}{dx} \left( \frac{2}{1 + \exp(Cx)} \right) \right|_{x=0} \quad (24)$$

obtaining

$$C = \frac{4}{\sqrt{\pi}} \quad (25)$$

Combining the result of Eqs. 25 and 23 with Eq. 22, we find the following approximation for the lower incomplete Gamma function:

$$Q\left(\frac{\mu^2}{\theta^2}, n\lambda \frac{\theta^2}{\mu}\right) \approx \frac{1}{1 + \exp\left(\sqrt{\frac{8}{\pi}} \frac{n\lambda - \mu}{\theta}\right)} \quad (26)$$

We rewrite terms of the form $(n\lambda \pm \bar{l})^a$ found before (Eq. 14) as

$$(n\lambda \pm \bar{l})^a = (n\lambda)^a \left(1 \pm \frac{\bar{l}}{n\lambda}\right)^a \quad (27)$$

and together with the approximations described above, and after appropriate simplifications aided by the Mathematica software, we find

$$\begin{aligned} v \approx & \frac{k_0 \theta (\mu^2 + \theta^2)}{\bar{l} \mu \sqrt{2\pi}} \sum_{n=1}^{\infty} \left[ \left(1 - \frac{\bar{l}}{n\lambda}\right)^{\mu^2/\theta^2} e^{\bar{l}\mu/\theta^2} + \left(1 + \frac{\bar{l}}{n\lambda}\right)^{\mu^2/\theta^2} e^{-\bar{l}\mu/\theta^2} - 2 \right] \left(\frac{n\lambda}{\mu}\right)^{\mu^2/\theta^2} e^{\frac{[\mu^2 - n\lambda\mu]}{\theta^2}} \\ & + \frac{2k_0 \mu^2}{\theta \sqrt{2\pi}} \sum_{n=1}^{\infty} \frac{\bar{l}}{n\lambda} \left(\frac{n\lambda}{\mu}\right)^{\mu^2/\theta^2} e^{\frac{[\mu^2 - n\lambda\mu]}{\theta^2}} \end{aligned} \quad (28)$$

The last term arises after simplification of the regularized Gamma terms approximated by the logistic function. Approximating $(1 \pm \bar{l}/\lambda)^k$ as an exponential:

$$(n\lambda + \bar{l})^k \approx (n\lambda + \bar{l})^k e^{\frac{k\bar{l}}{n\lambda}} \quad (29)$$

one obtains

$$\begin{aligned} v \approx & \frac{k_0}{\sqrt{2\pi}} \frac{\theta(\mu^2 + \theta^2)}{\mu \bar{l}} \sum_{n=1}^{\infty} \left( e^{\frac{\mu(n\lambda - \mu)\bar{l}}{\theta^2 n\lambda}} + e^{\frac{\mu(\mu - n\lambda)\bar{l}}{\theta^2 n\lambda}} - 2 \right) e^{\frac{(\mu - n\lambda)\mu}{\theta^2}} \\ & + \frac{2k_0 \mu^2}{\theta \sqrt{2\pi}} \sum_{n=1}^{\infty} \frac{\bar{l}}{n\lambda} \left(\frac{n\lambda}{\mu}\right)^{\mu^2/\theta^2} e^{\frac{(\mu - n\lambda)\mu}{\theta^2}} \end{aligned} \quad (30)$$

The velocities as a function of the mean jump length and the noise are shown in Figure S2A. The first term with the exponentials in the parenthesis in Eq. 30 is responsible for growing and decaying from $\mu = n\lambda$, Figure S2B: when $n\mu$ is a multiple of $\lambda$ both exponential terms are 1, and together with the adjacent factor 2, results in the full cancellation of the first term. On the other hand, the last term defines the main response at $\mu = n\lambda$ (Figure S2C). Based on these observations, from now on, we will use

$$v \approx \frac{2k_0 \mu^2}{\theta \sqrt{2\pi}} \sum_{n=1}^{\infty} \frac{\bar{l}}{n\lambda} \left(\frac{n\lambda}{\mu}\right)^{\mu^2/\theta^2} e^{\frac{(\mu - n\lambda)\mu}{\theta^2}}, \quad (31)$$

as the simplest possible equation describing the velocity of the motor. Although this expression significantly overestimates the velocity, it remains useful for analytically describing changes of the velocity with the parameters, as it will become evident in the following sections.

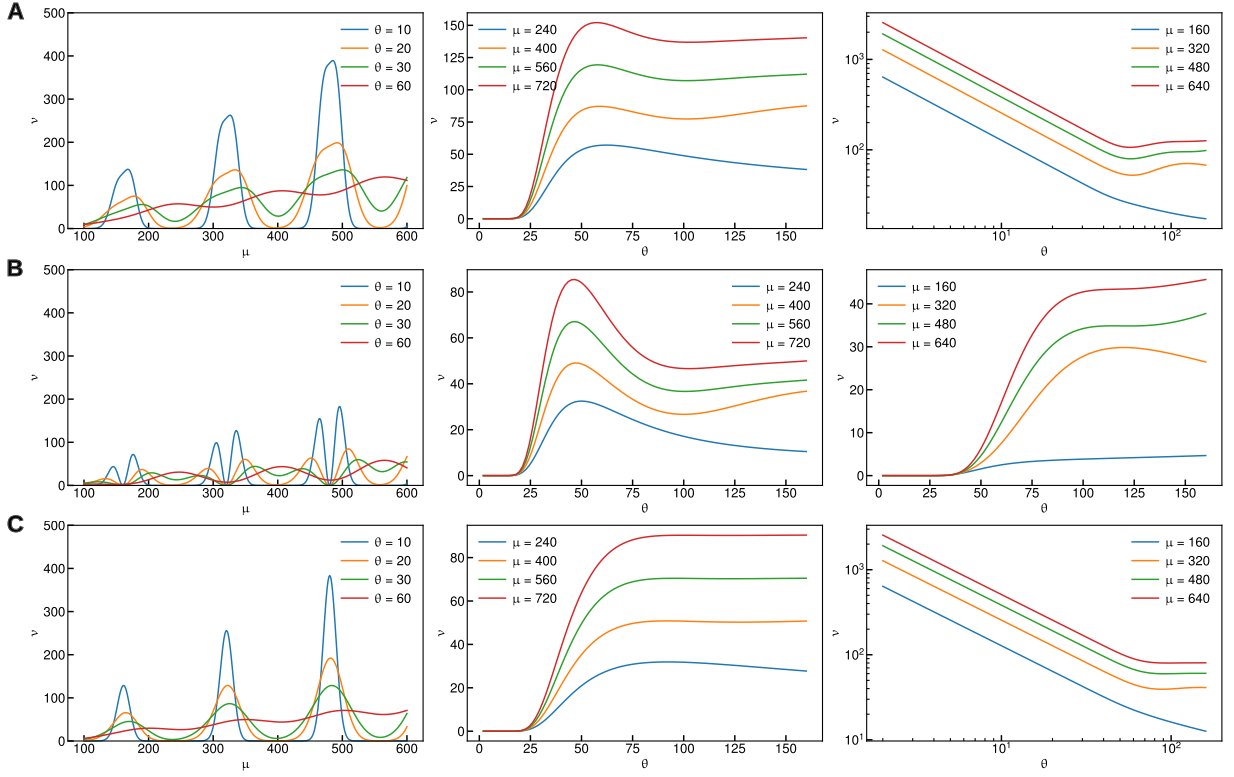

**Figure S2: Approximation of the heuristic model in terms of elementary functions.** Velocity as a function of the mean jump length and as a function of noise (in the anti-phase and in-phase case). (a) Full expression as given by Eq. 30. (b) Velocity given by the first term in Eq. 30. (c) Velocity in simplest functional form Eq. 31.

*Dependency of the optimal mean jump on the noise:* First, we seek to understand the change of the peak velocity for fixed $\theta$. The numerical model and the theory indicate that for zero variance, the optimum step is $\mu_0^* = \lambda$. To determine how $\mu^*$ depends on $\theta$, we analyze the leading order term of the summation $n = 1$, and we look for the maximum

$$\left. \frac{\partial v(\mu, \theta)}{\partial \mu} \right|_{\theta} \stackrel{!}{=} 0 \quad (32)$$

resulting in

$$\frac{\partial v(\mu, \theta)}{\partial \mu} = \frac{k_0 \bar{l}}{\theta^3} \sqrt{\frac{2}{\pi}} e^{-\frac{\mu(\lambda - \mu)}{\theta^2}} \left( \frac{\lambda}{\mu} \right)^{\frac{\mu^2}{\theta^2} - 1} \left[ 2\mu^2 \log \left( \frac{\lambda}{\mu} \right) - \mu(\lambda - \mu) + 2\theta^2 \right] \stackrel{!}{=} 0 \quad (33)$$

The term in the parenthesis has to be zero to satisfy the equality. Since for $\theta \ll \mu$ the optimal step is located at $\mu^* = \lambda$, we Taylor expand the logarithm up to second order around $\lambda/\mu \equiv 1$:

$$\log \left( \frac{\lambda}{\mu} \right) \approx \left( \frac{\lambda}{\mu} - 1 \right) - \frac{1}{2} \left( \frac{\lambda}{\mu} - 1 \right)^2 \quad (34)$$

and insert this into the term in the parenthesis above, gives a second order polynomial equation for $\mu$ which solution yields:

$$\mu^* \approx \frac{\lambda}{4} \left[ 3 + \sqrt{1 + \left( \frac{2\theta}{\lambda} \right)^2} \right]. \quad (35)$$

Although this has been developed for the first obstacle, we expect that the approach should work for any $\mu_0 \approx c\lambda$, where $c \in \mathbb{N}$ by appropriately rescaling the variables (Figure S4).

Alternatively, we can pose the ansatz that the optimal velocity occurs when the peak of the step size matches $\lambda$. The optimal of the Gamma distribution is $(a-1)b^{-1}$. Then, in terms of the moments, we look for

$$\mu - \frac{\theta^2}{\mu} \stackrel{!}{=} \lambda \quad (36)$$

which after solving for $\mu$ yields:

$$\mu^* = \frac{\lambda}{2} \left[ 1 + \sqrt{1 + \left( \frac{2\theta}{\lambda} \right)^2} \right]. \quad (37)$$

Note the similarity between the functional form obtained and that found from the heuristic model. Moreover, both approaches predict that for $\theta/\mu \rightarrow 0$, the critical $\mu^*$ coincides with the wavelength of the system, $\mu^* \rightarrow \lambda$.

By rescaling the $\vartheta := \theta^2/(\mu_0\lambda)$ and the critical $\bar{\mu}^* = \mu^*/\mu_0$ by the mean jump length for the successive in-phase jump $\mu_0 := n\lambda$, we find master curves for $n > 1$ (Figure S4). Interestingly, the ansatz method has good predictive power for the small noise regime for $n = 1$, i.e., $\mu_0 = \lambda$. However, for the successive multiples, the predicted critical mean jump is better described by the approximated formula derived from the heuristic model (Eq. 35). Both formulas underestimate the critical mean jump under high noise conditions.

*Dependency of the velocity on the noise (in-phase case):* Next, we determined the dependency on $\theta$ for a fixed step size. First, we study the favorable in-phase case, where $\mu \approx \lambda$. Then, we set $\mu = \lambda$ in Eq. 31, and for $n = 1$,

$$v_1 = \frac{2k_0\lambda\bar{l}}{\theta\sqrt{2\pi}} \quad (38)$$

The full equation is:

$$v = v_1 + \frac{2k_0\lambda\bar{l}}{\theta\sqrt{2\pi}} \sum_{n=2}^{\infty} n^{\frac{\lambda^2}{\theta^2}-1} e^{\frac{\lambda^2(1-n)}{\theta^2}} \quad (39)$$

or if we use the dimensionless term $\bar{\theta} := \theta/\lambda$,

$$\begin{aligned} v &\approx v_1 + \frac{2k_0\bar{l}}{\bar{\theta}\sqrt{2\pi}} \sum_{n=2}^{\infty} n^{\frac{1}{\bar{\theta}^2}-1} e^{\frac{1-n}{\bar{\theta}^2}} \\ &\approx v_1 + \frac{2k_0\bar{l}}{\bar{\theta}\sqrt{2\pi}} \sum_{n=2}^{\infty} e^{\frac{(1-n)+(\bar{\theta}^2-1)\log n}{\bar{\theta}^2}} \end{aligned} \quad (40)$$

For the range of values investigated in this work $\bar{\theta} < 1$, and the exponential term in the summation is then always negative. Moreover, the contribution of the sum term decays quickly with the $n$ terms and with decreasing $\theta$ due to the $\theta^{-2}$ term. We conclude that velocity for small $\theta$ in the in-phase case is thus dominated by $v_1$, and therefore the velocity scales as $v \sim \theta^{-1}$.

We show the comparison between the simulation and the theoretical predictions in Figure S5A. We rescaled the velocities by their corresponding mean jump and the standard deviation by the linker dimensions, finding master curves for the full regime. The power law decay shows excellent agreement over a wide range of noise levels. Interestingly, however, for the very low noise level $\theta < \bar{l}/2$ the simulations show a lower exponent, which is consistent with the motor having a negligible probability of hitting the obstacle during jumps.

*Dependency of the velocity on the noise (anti-phase case):* In the unfavorable anti-phase scenario, the step size $\mu$ is thus making the motor to jump, on average, on the middle of the obstacle:

$$\mu_u = \frac{2n+1}{2}\lambda, \quad (41)$$

thus impairing the progress of the motor. For small $\theta/\lambda$ the motor velocity is zero, since it does not find available space. Nevertheless, we observe that upon increasing $\theta$ there is a transition that allows the motor to progress. This can be identified by the sigmoidal-like shape of $v$ as a function of $\theta$ (Figure S3C). We seek to determine a characteristic $\theta^*$ at which this transition occurs. The first instinct is to solve for the inflection point of the curve. Still, due to the complicated functional form of Eq. 31, the second derivative becomes extremely complex, making an analytical solution unlikely. Therefore, we will employ the half-maximum velocity, $v_{\text{HM}}$, as a proxy for the transition. We restrict the analysis to the first term, $n = 1$:

$$v \approx \frac{2k_0\mu^2}{\theta\sqrt{2\pi}} \frac{\bar{l}}{\lambda} \left(\frac{\lambda}{\mu}\right)^{\mu^2/\theta^2} e^{\frac{(\mu-\lambda)\mu}{\theta^2}} \quad (42)$$

Now, we define $\mu = u\lambda$, where $u \in (1, 3/2)$, so that $\mu_u = 3\lambda/2$ represents the completely anti-phase case for the first couple of obstacles. The maximum velocity $v_{\text{max}}$ can be determined by setting $\theta = \lambda/2$, where the factor $1/2$ is a mathematically convenient term that phenomenologically captures the value of $\theta$ resulting in the maximum velocity. Then, $v_{\text{HM}} = v_{\text{max}}/2$ can be written as:

$$v_{\text{HM}} = \frac{2k_0\bar{l}u^2\lambda}{\sqrt{2\pi}} \left(\frac{1}{u}\right)^{4u^2} e^{4u(u-1)} \quad (43)$$

We look for the critical $\theta^*$ by solving the equation $v \stackrel{!}{=} v_{\text{HM}}$:

$$\frac{2k_0\bar{l}u^2\lambda}{\theta\sqrt{2\pi}} \left(\frac{1}{u}\right)^{u^2\lambda^2/\theta^2} e^{\frac{\lambda^2(u-\lambda)u}{\theta^2}} \stackrel{!}{=} v_{\text{HM}}. \quad (44)$$

By introducing the dimensionless noise $\bar{\theta} := \theta/\lambda$, the equation to solve is simplified to

$$\frac{1}{\bar{\theta}} \left(\frac{1}{u}\right)^{\frac{u^2}{\bar{\theta}^2}} e^{\frac{(u-1)u}{\bar{\theta}^2}} = \left(\frac{1}{u}\right)^{4u^2} e^{4u(u-1)}. \quad (45)$$

Expressing the power in terms of an exponential, we obtain:

$$\frac{1}{\bar{\theta}} \exp\left(\frac{u^2(1 - \log u) - u}{\bar{\theta}^2}\right) = \exp(4u^2(1 - \log u) - 4u) \quad (46)$$

and after taking logarithms and regrouping terms, we find the transcendental equation:

$$-\log \bar{\theta} + \frac{u^2(1 - \log u) - u}{\bar{\theta}^2} = 4u^2(1 - \log u) - 4u. \quad (47)$$

After expansion of the logarithm around $\bar{\theta} = 1/2$, which corresponds to the position of maximum velocity, we obtain:

$$\log \bar{\theta} \approx \log 2 + 2\left(\bar{\theta} - \frac{1}{2}\right), \quad (48)$$

one finds the following cubic equation

$$2\bar{\theta}^3 + (B - 1 - \log 2)\bar{\theta}^2 - A = 0. \quad (49)$$

where $A = u^2(1 - \log u) - u$ and $B = \log 2 + 4u^2(1 - \log u) - 4u$. To find the roots, we rewrite it in depressed form:

$$t^3 + pt + q = 0 \quad (50)$$

where

$$t \equiv \bar{\theta} + \frac{1}{6}(B - 1 - \log 2), \quad (51)$$

and

$$p \equiv -\frac{1}{12}(B - 1 - \log 2)^2, \quad q \equiv \frac{1}{216} [2(B - 1 - \log 2)^3 - 108A] \quad (52)$$

The discriminant $\Delta = -(4p^3 + 27q^2)$ is positive, and then the equation admits three real solutions. Hence, Viète's trigonometric method can be used to express the roots of the depressed cubic equation:

$$t_m = 2\sqrt{-\frac{p}{3}} \cos \left[ \frac{1}{3} \cos^{-1} \left( \frac{3q}{2p} \sqrt{-\frac{3}{p}} \right) - \frac{2\pi m}{3} \right], \quad (53)$$

where $m = \{0, 1, 2\}$. In particular, we find that, upon transforming back $t$ to $\bar{\theta}$ (Eq 51), the results for $m = 1$ are consistent with the values observed in the simulations (Figure S5B). Since the result function is quasi-linear, we expand $\theta^*$ up to first order around $u = 5/4$, the middle value between being completely in-phase or completely anti-phase, obtaining

$$\bar{\theta} \approx \theta_0 + \theta_1 \left( u - \frac{5}{4} \right), \quad (54)$$

where $\theta_1 \approx 0.153719$ and $\theta_2 \approx 0.658647$ are the numerical values obtained from the evaluation of the function and its derivative.

We find good agreement between theoretical predictions and the simulation results for every $\mu_u$, with the larger discrepancy occurring near the completely anti-phase case, $u = 3/2$ (Figure S5B). In our analyses, for theoretical tractability, we assumed symmetry with respect to $\mu_u$, such that $u = |\mu - \mu_u| \lambda^{-1}$. Simulations show however that for a given mean jump length between obstacles, the characteristic $\theta$ is higher for $\mu < \mu_u$. The difference decreases as $u$ approximates 1 or $3/2$, as well as with the increase in the signal-to-noise ratio, $\mu_u/\theta$.

#### B.2. Numerical evaluation of the model

We employed Python to obtain numerical values from Eq. 12 for representation. We used the functions `gamma` and `gammainc` of the SciPy library. However, for some couples of values $\mu - \theta^2$, the associated $b$ parameter is very small, making the upper incomplete Gamma functions $\Gamma(a, x)$ present in the calculation of the velocity to overflow. To overcome this limitation, we used the `mpmath` library, which allows to perform arbitrary-precision floating-point arithmetic. Nevertheless, even by using the `mpmath` library, some values can still be numerically unstable or require long calculation times. In such cases, we use the approximation of the incomplete Gamma function $Q$ for small $b$ in terms of the complementary error function (Eq. 22).

The summation in Eq. 12 is truncated to $n_t$ terms by accounting for the maximum $\mu$ analyzed, such that the probability of the motor jumping the full trail of obstacles is negligible, i.e., $n_t \gg \mu/\lambda$. We truncated the summation to $n_t = 20$ for the pseudo-analytical calculations shown in this work.

### C. Link between fat-tail waiting time distributions and sub-linear average trajectories

To understand the relationship between the distributions of waiting times between jumps and the shape of the average trajectory of the position over time, let us consider a model simpler than our jump model. Suppose that the hinge performs jumps of constant step size $\langle \Delta x \rangle$, but with waiting times $\Delta t$ between consecutive jumps that follow a distribution $p(\Delta t)$. This is a classic case of a renewal process [4]. The probability $P(n, t)$ of being at position $n\langle \Delta x \rangle$ at time $t$ is given by the system of implicit equations:

$$P(0, t) = 1 - \int_0^t d\Delta t p(\Delta t) \quad (55)$$

$$P(n > 0, t) = \int_0^t d\Delta t P(n-1, t-\Delta t) p(\Delta t) \quad (56)$$

For the simple case of an exponential distribution of waiting times ($p(\Delta t) = k_0 e^{-k_0 \Delta t}$) corresponding to a case without obstacles in our framework, we can easily show by induction that $P(n, t)$ follows a Poisson distribution of rate parameter $k_0 t$ ($P(n, t) = (k_0 t)^n / (n!) e^{-k_0 t}$). Thus:

$$\langle x(t) \rangle \equiv \langle \Delta x \rangle \sum_{n=0}^{\infty} n P(n, t) = \langle \Delta x \rangle k_0 t \quad (57)$$

Now assume that the waiting time distribution has the form $p(\Delta t) = b(a-1)/(1+b\Delta t)^a$ with $a > 1$ to ensure that $\int_0^{\infty} d\Delta t p(\Delta t)$ is finite. If $a \leq 2$, the mean of $p(\Delta t)$ tends towards infinity and we speak of a 'fat-tail' distribution. For $n = 0$, we can easily calculate the integral Eq. 55 giving $P(n=0, t) = 1/(1+bt)^{a-1}$. To solve Eq. 56, we notice that $P(n, t)$ is the convolve of $P(n-1, t)$ with $p(\Delta t)$, therefore the Laplace transform of $P(n, t)$ by definition equal to $\tilde{P}(n, s) \equiv \int_0^{\infty} dt P(n, t) e^{-st} = \tilde{P}(n-1, s) \tilde{p}(s)$, or $\tilde{P}(n, s) = \tilde{p}^n(s) \tilde{P}(0, s)$. Now, by definition of $P(0, t)$ (Eq. 55), $\tilde{P}(0, s) = (1 - \tilde{p}(s))/s$, therefore:

$$\tilde{P}(n, s) = \frac{\tilde{p}^n(s)(1 - \tilde{p}(s))}{s} \quad (58)$$

Thus,

$$\begin{aligned}\langle \tilde{x} \rangle(s) / \langle \Delta x \rangle &= \sum_n n \tilde{P}(n, s) = \sum_n n \frac{\tilde{p}^n(s)(1 - \tilde{p}(s))}{s} \\ &= \left( \frac{1 - \tilde{p}(s)}{s} \right) \left( \frac{\tilde{p}(s)}{(1 - \tilde{p}(s))^2} \right) = \frac{\tilde{p}(s)}{s(1 - \tilde{p}(s))}\end{aligned}\quad (59)$$

It can be shown that

$$\tilde{p}(s) = (a - 1)e^{s/b}(s/b)^{a-1}\Gamma_i[1 - a, s/b] \quad (60)$$

where $\Gamma_i[v, x] \equiv \int_x^\infty dt t^{v-1}e^{-t}$ is the Euler's upper incomplete gamma function. As $s \rightarrow 0$,

$$\tilde{p}(s) \sim 1 - \Gamma[2 - a](s/b)^{a-1} \quad \text{for } 1 < a < 2 \quad (61)$$

$$\sim 1 - s/(b(a - 2)) \quad \text{for } a > 2 \quad (62)$$

where $\Gamma[v] \equiv \Gamma_i[v, 0]$ is Euler's gamma function. Substituting this back into Eq. 59, we find that as $s \rightarrow 0$

$$\langle \tilde{x} \rangle(s) / \langle \Delta x \rangle \sim \frac{1}{b\Gamma[2 - a]} \left( \frac{b}{s} \right)^a \quad \text{for } 1 < a < 2 \quad (63)$$

$$\sim \frac{b(a - 2)}{s^2} \quad \text{for } a > 2 \quad (64)$$

Inverting the transform, in the limit $t \rightarrow \infty$ (dominated by the $s \rightarrow 0$ limit), we obtain

$$\langle x \rangle(t) / \langle \Delta x \rangle \sim (bt)^{a-1} / (\Gamma[2 - a]\Gamma[a]) \quad \text{for } 1 < a < 2 \quad (65)$$

$$\sim (a - 2)(bt) \quad \text{for } a > 2 \quad (66)$$

Thus, in the fat-tail case ($1 < a < 2$), we obtain sub-linear average progression, whereas the progression is linear when the expected waiting time is finite ($a > 2$).

### III. OTHER SUPPLEMENTARY FIGURES

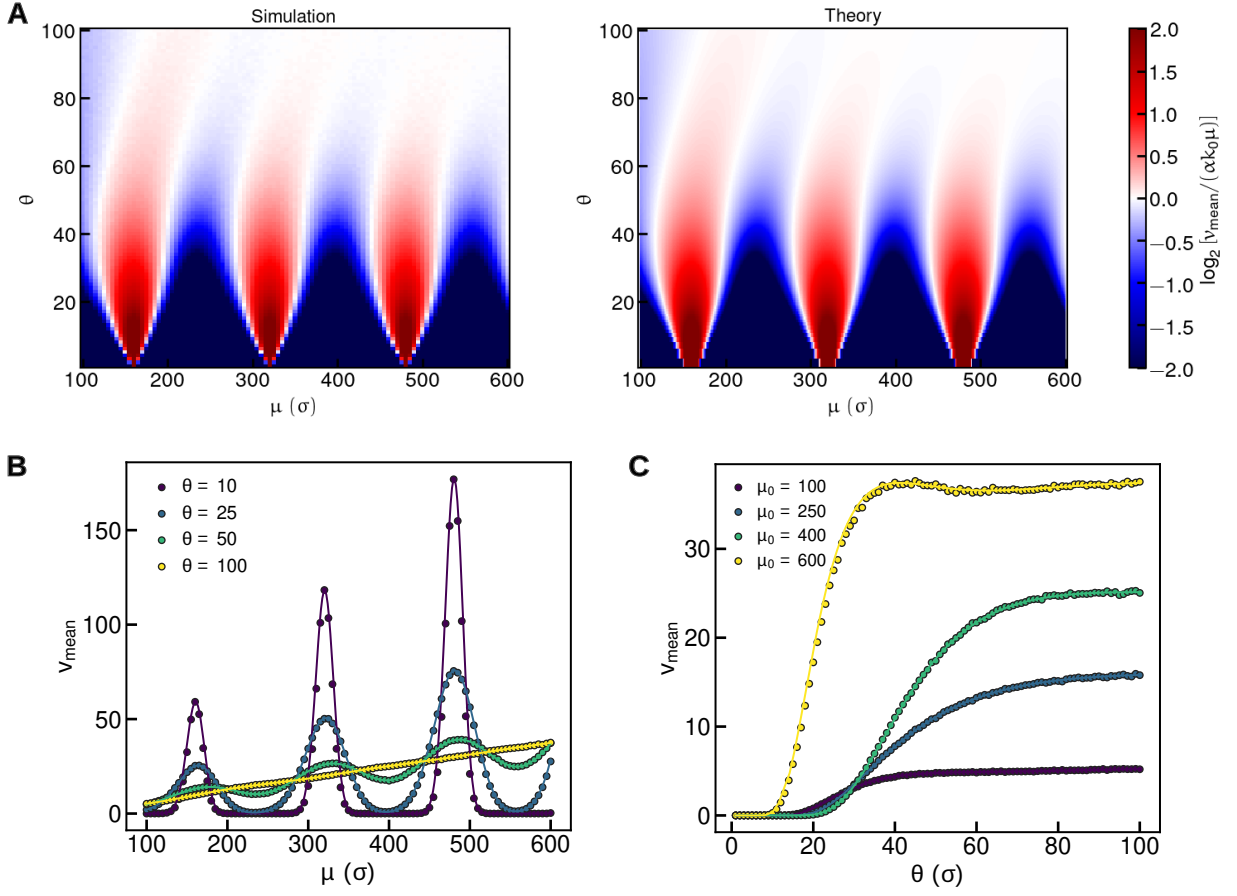

**Figure S3: Velocity of the motor as a function of the jump distribution parameters. Comparison between simulation and theory.** (a) General overview of the velocities obtained in the simulation (right) and the theoretical predictions (left) for the same pairs of parameters $\mu$ and $\theta$. (b) Velocity of the motor as a function of the mean step $\mu$ for fixed noise level, $\theta$. (c) Velocity of the motor as a function of the noise for fixed mean step. The dots are the result of the simulation and the continuous lines are the heuristic model predictions. Results for $\bar{l} = 10$ and $s = 150$.

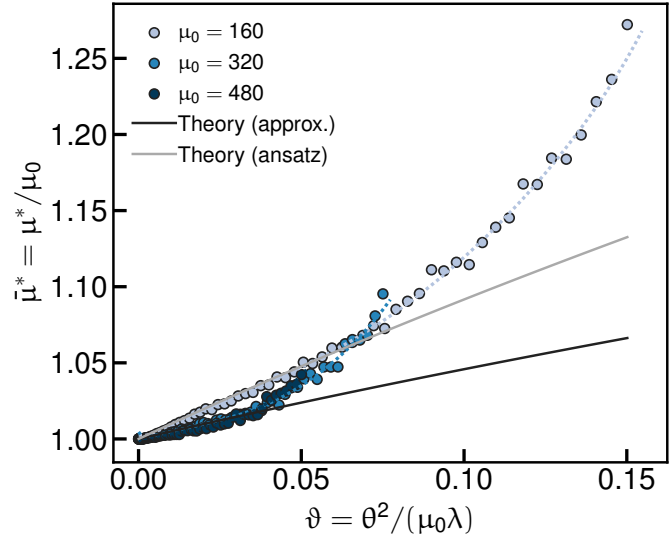

**Figure S4: Critical mean jump giving the maximum velocity as a function of the dispersion level.** Result for fixed mean step. The dots are the result of the simulation and the dotted lines are the predictions of the full heuristic model. Predictions of the simplified heuristic model (Eq. 35) and from the ansatz formula (Eq. 37) are shown with continuous lines. Results for linker lengths of $\bar{l} = 10$ and obstacle dimensions $s = 150$.

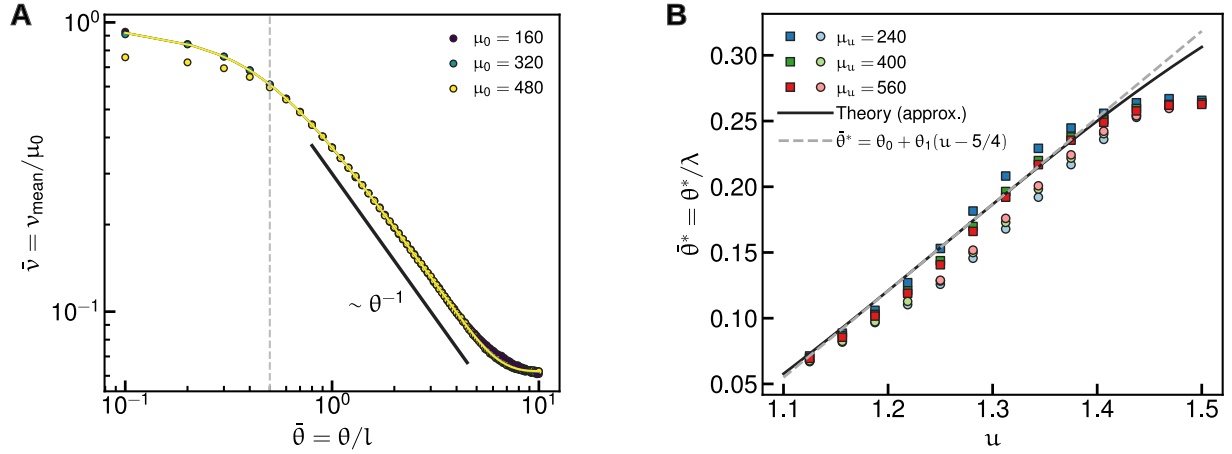

**Figure S5: Velocity changes with dispersion for the in-phase and anti-phase scenarios.** (a) Decay of the velocity with the variability of the motor jumps $\theta$ in the in-phase case. The dots are the results of the simulation and the continuous lines are the results of heuristic model. The thick black line shows the predicted scaling from Eq. 38. The dotted vertical line highlight the change of trend for $\bar{l}/2$. (b) Critical $\theta$ value resulting in half-maximum velocity in the anti-phase scenario. The square symbols in dark colors represent simulation results for $\mu < \mu_u$; circles in light pastel colors are their counterpart for $\mu > \mu_u$. The black line is the closed analytical solution. The dashed lined in gray is the linear approximation. Data for $\bar{l} = 10$ and $s = 150$.

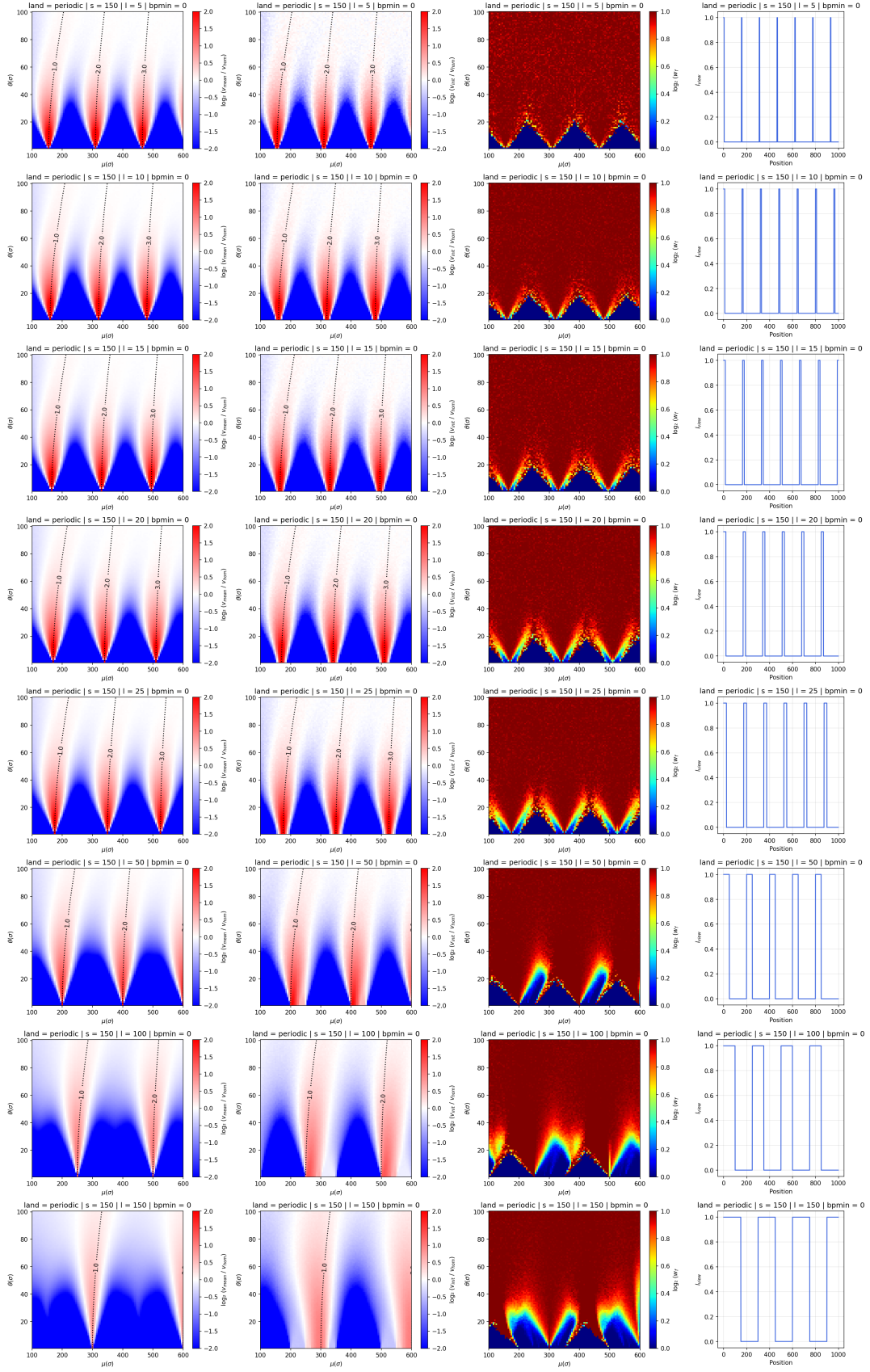

**Figure S6: Velocity in periodic landscapes: dependency on linker length.** Linker lengths $\bar{l}$ in rows, increase from top to bottom. In columns, from left to right: $\log_2(\frac{v_{\text{mean}}}{v_{\text{hom}}})$, $\log_2(\frac{v_{\text{init}}}{v_{\text{hom}}})$, $w_f$, and typical profiles of accessibility. Data for $s = 150$.

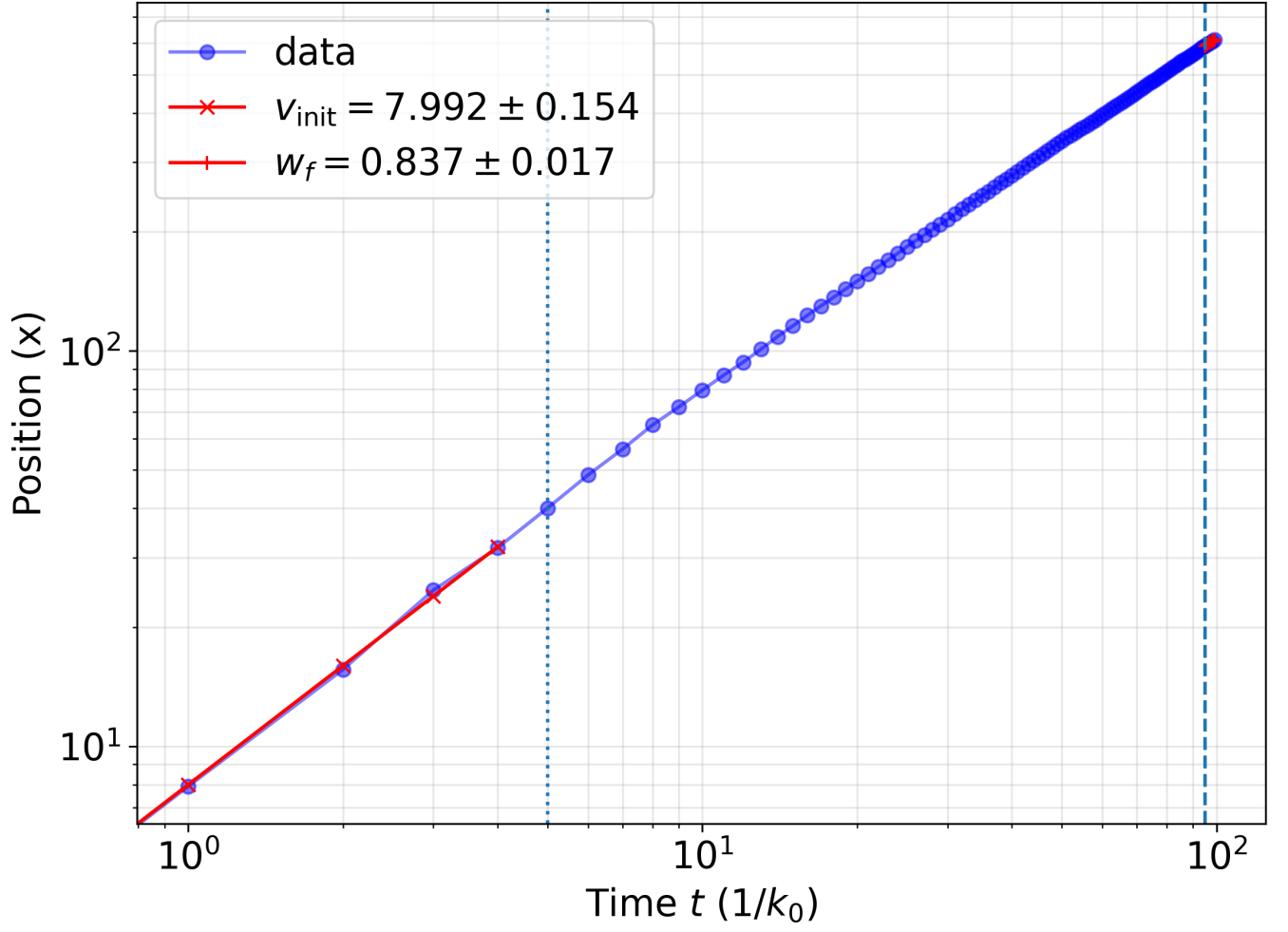

**Figure S7: Determination of the initial velocity and the exponent $w_f$ in random landscapes.** The initial velocity $v_{\text{init}}$ is determined from a linear fit over the time window $t = [0, 5/k_0]$ (dotted vertical line). The late-time scaling $w_f$ is determined from fitting the position to a function of the form $\langle x(t) - x(0) \rangle = Ct^{w_f}$ for $t \geq 0.8t_{\text{max}}$ (dashed vertical line). Data for a random scenario ($\bar{l} = 10$, $s = 150$) with $\mu = 150$ and $\theta = 100$.

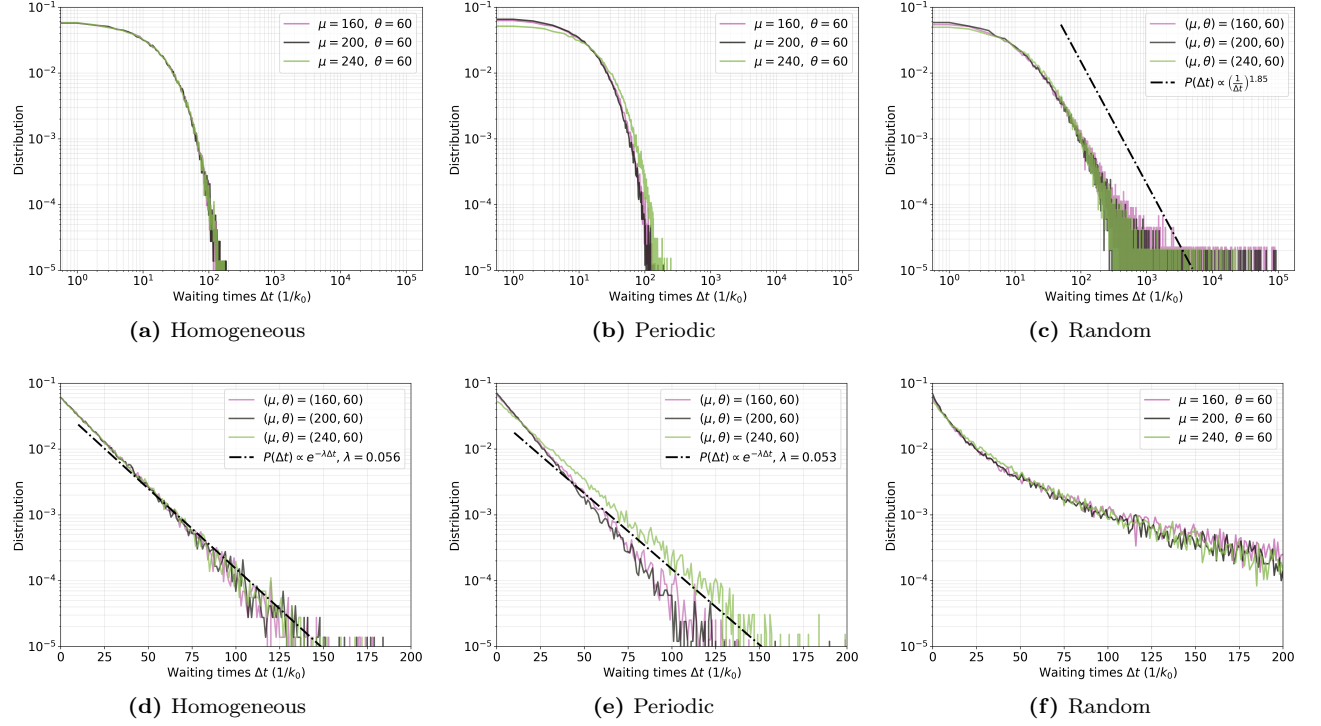

**Figure S8: Distributions of waiting times.** Comparison of the distribution of waiting times for different $\mu$ and $\theta$ for homogeneous (left), periodic (center) and random (right) landscapes. Top row shows the data in log-log scale.

Bottom row, same data in log scale. While homogeneous and periodic scenarios always exhibit exponential distributions, random case may lead to scaling-law, fat-tail distributions. Data for $\bar{l} = 10$ and $s = 150$.

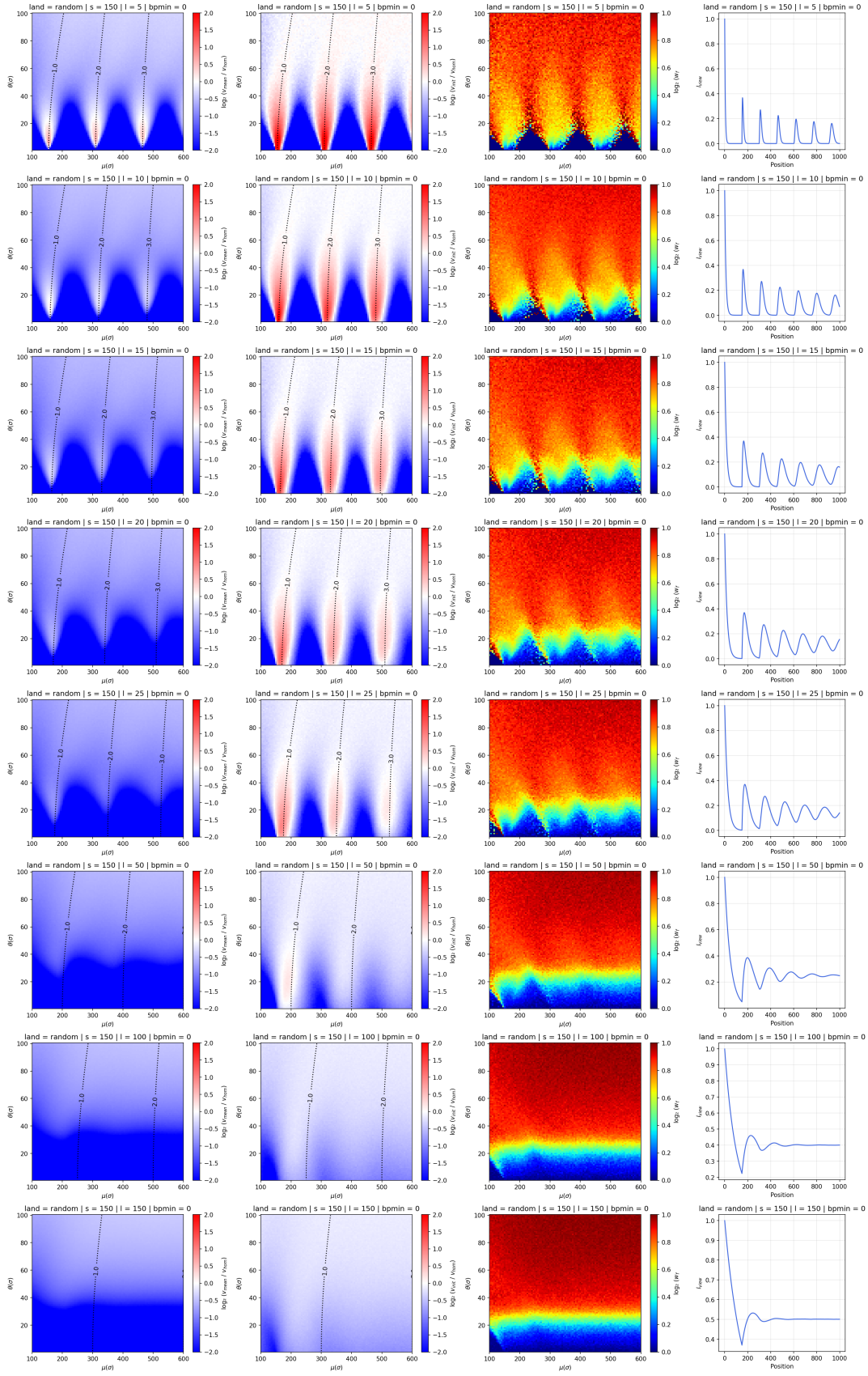

**Figure S9: Velocity in random landscapes: dependency on linker length.** Linker lengths $\bar{l}$ in rows, increase from top to bottom. In columns, from left to right: $\log_2(\frac{v_{\text{mean}}}{v_{\text{hom}}})$, $\log_2(\frac{v_{\text{init}}}{v_{\text{hom}}})$, $w_f$, and average accessibility profile after an obstacle. Data for $s = 150$.

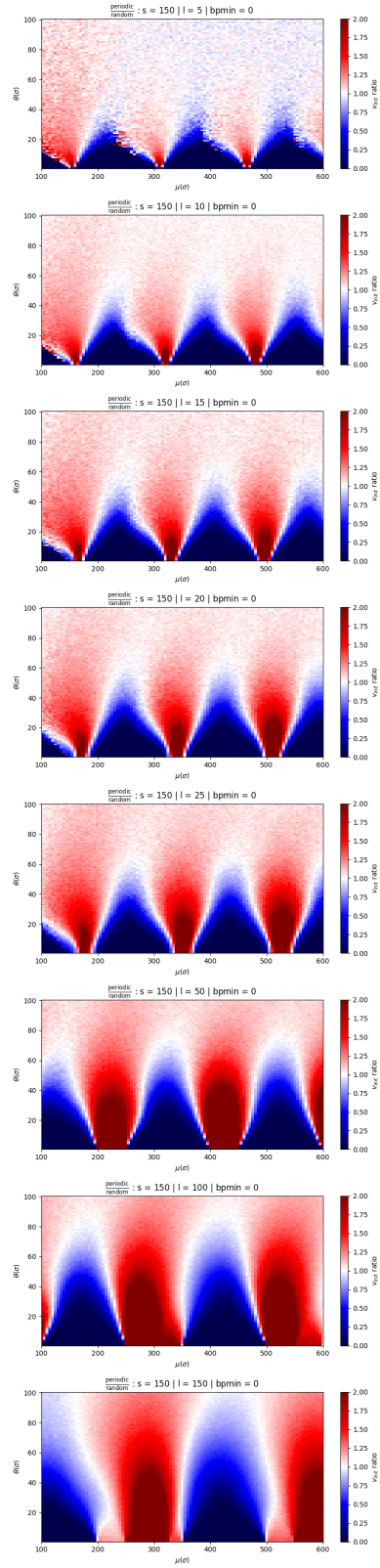

**Figure S10: Comparison of velocities between the periodic and random landscapes.** The ratio $v_{\text{init}}^{\text{periodic}}/v_{\text{init}}^{\text{random}}$ is shown for various linker values $\bar{l}$ ($\bar{l}$ increases from top to bottom). Extrusion in the periodic scenario is faster than in the random case in 'in-phase' regimes and slower in 'anti-phase' regimes. Data for $s = 150$.

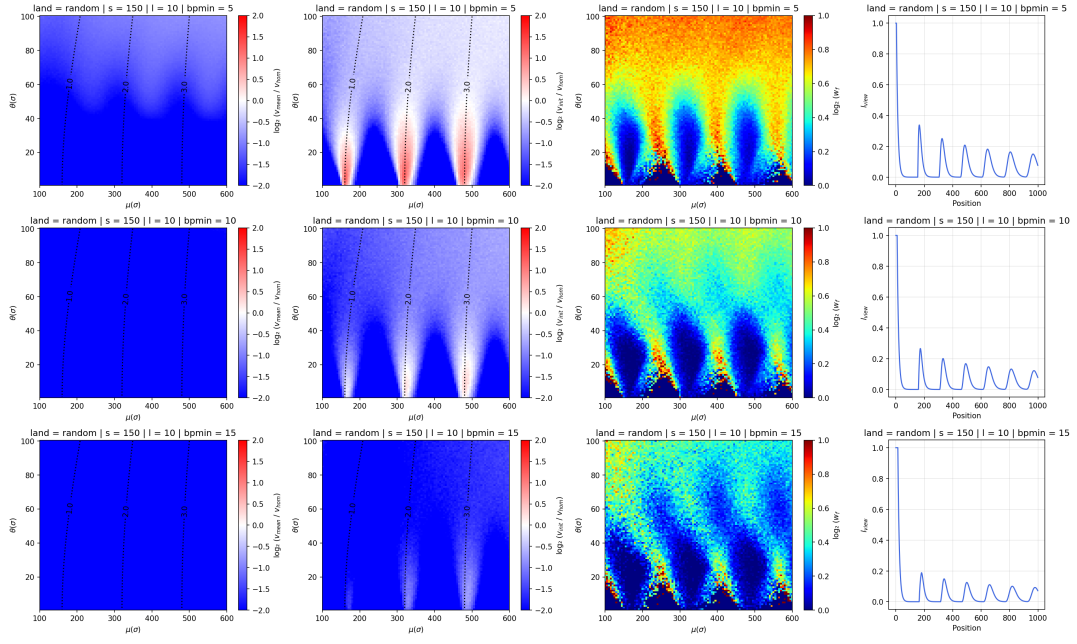

**Figure S11: Velocity in random landscapes: dependency on the minimal binding size.** Minimal binding site $bp_{min}$ in rows, increase from top to bottom. In columns, from left to right: $\log_2(\frac{v_{mean}}{v_{hom}})$, $\log_2(\frac{v_{init}}{v_{hom}})$, $w_f$, and average accessibility profile after an obstacle. Data for $\bar{l} = 10$ and $s = 150$.

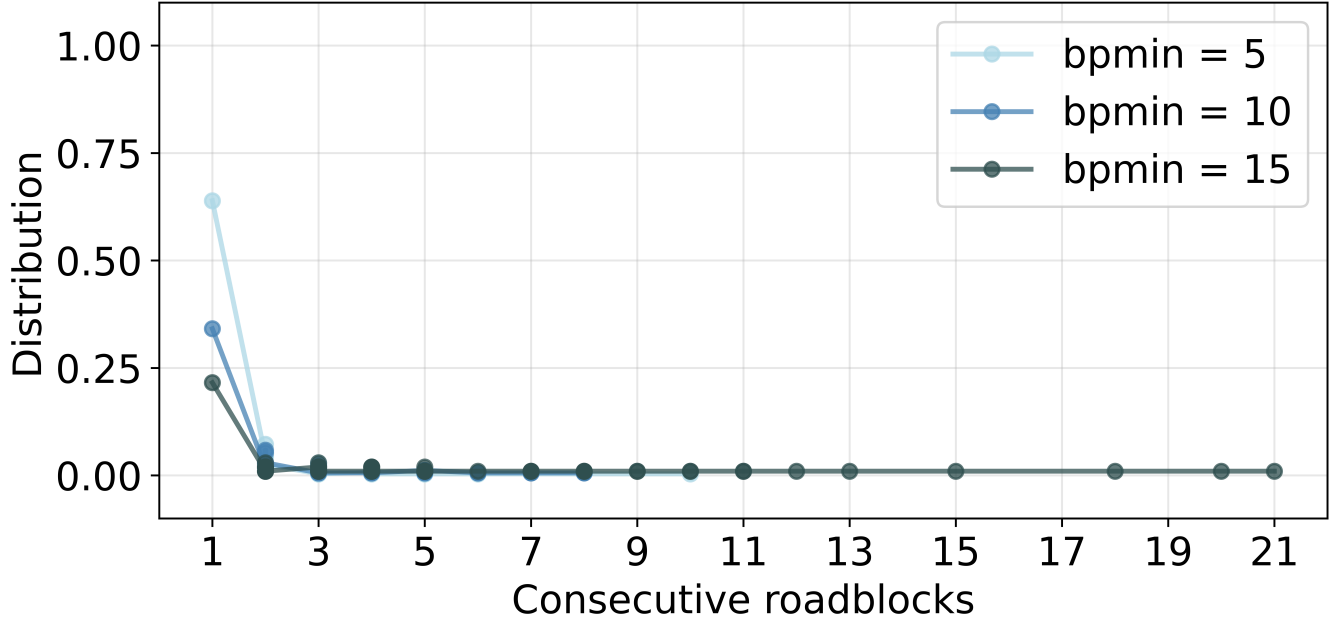

**Figure S12: Consecutive roadblocks in presence of a minimal binding size.** Distribution of the number of roadblocks present effective obstacle in for different minimal binding size $bp_{min}$. Increasing $bp_{min}$ leads to longer obstacles made of several roadblocks. Data for $\bar{l} = 10$ and $s = 150$.

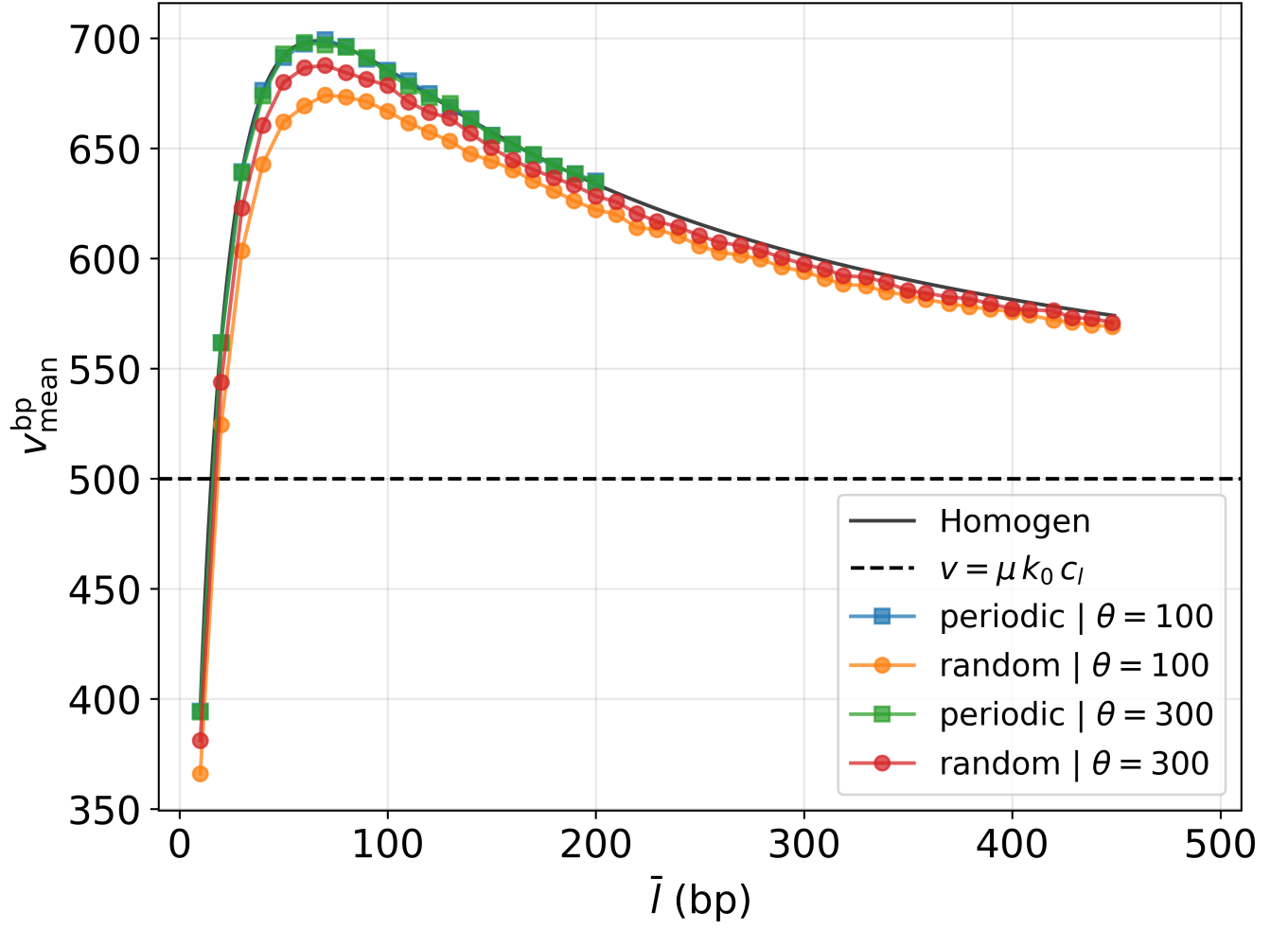

**Figure S13: Impact of chromatin compaction on loop extrusion speed for $\mu = 500$.** Dependency of the mean bp-velocity $v_{\text{mean}}^{\text{bp}}$ (in $k_0 \text{bp}$ units) on linker length $\bar{l}$. Results for different landscapes and values of $\theta$ for fixed mean step size $\mu = 500$ ($s = 35$).

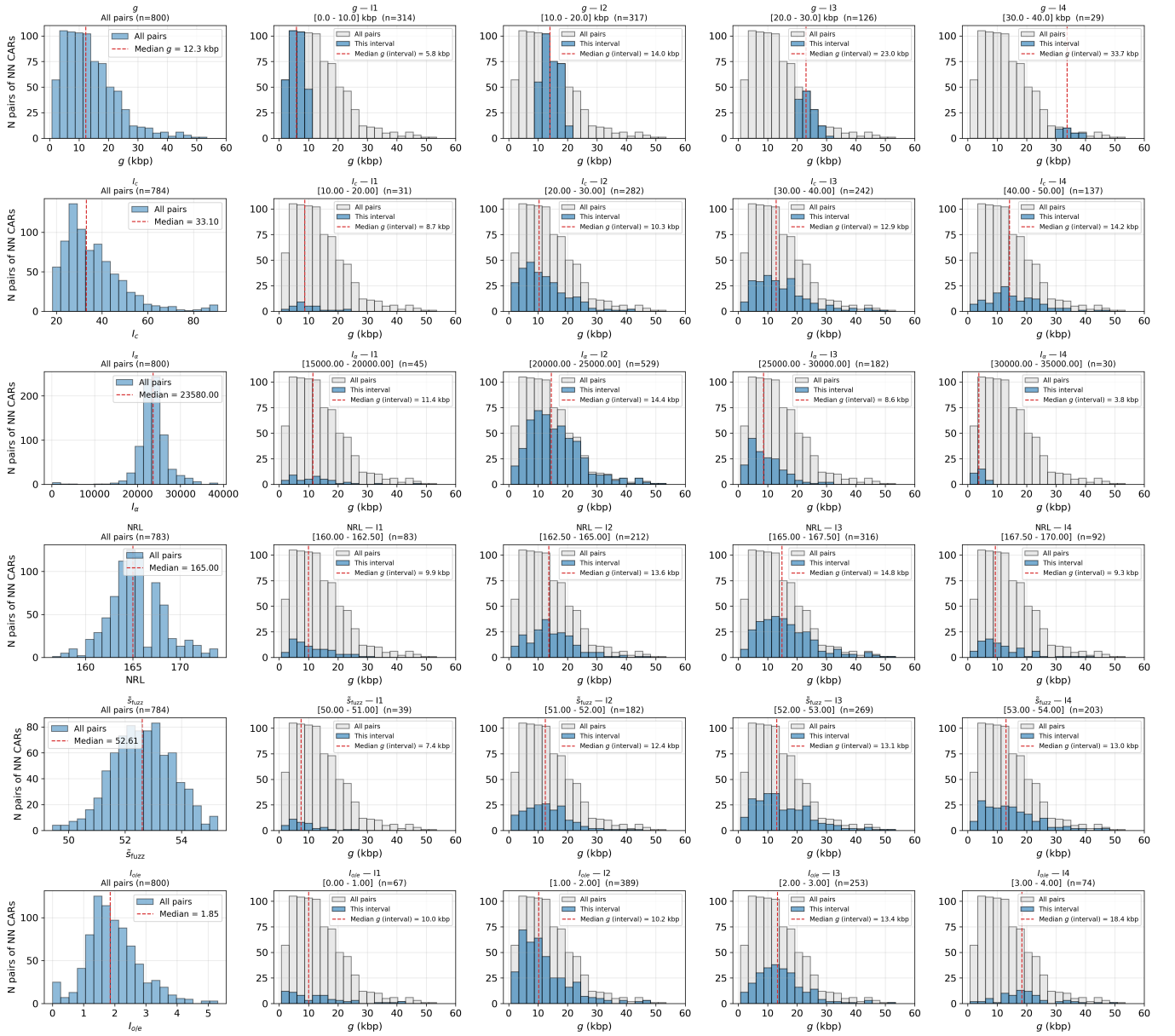

**Figure S14: Properties of nucleosome landscapes between CARs.** Panels in the most left column are the distributions of quantities characterizing the nucleosome landscape between consecutive CARs: (from top to bottom) the genomic distance $g$ between CARs, the intensity $I_c$ of cohesin ChIP-seq signal around CARs, the median nucleosome density $I_\alpha$ between the CARs, the median $\widetilde{NRL}$ between the CARs, the median fuzziness score $\tilde{s}_{fuzz}$ between the CARs and the contact enrichment $I_{o/e}$ between the CARs. For each metrics, four equal-length intervals ($I_1$ to $I_4$) are defined covering the full range of values. Columns 2 to 5 represents the distribution of genomic distances between the pairs of consecutive CARs belonging to each interval. $g$ has a fairly wide distribution typically varying between 1 and 40 kbp with a median value at 12.5 kbp. $I_c$ also has a broad distribution and is positively correlated with genomic distance. $I_\alpha$ has a pitted distribution and does not appear to have a clear correlation with $g$. $\widetilde{NRL}$ and $\tilde{s}_{fuzz}$ are sparsely dispersed, have bell-shaped distributions centered around 165 bp and 52.6 respectively, and are poorly correlated with $g$.

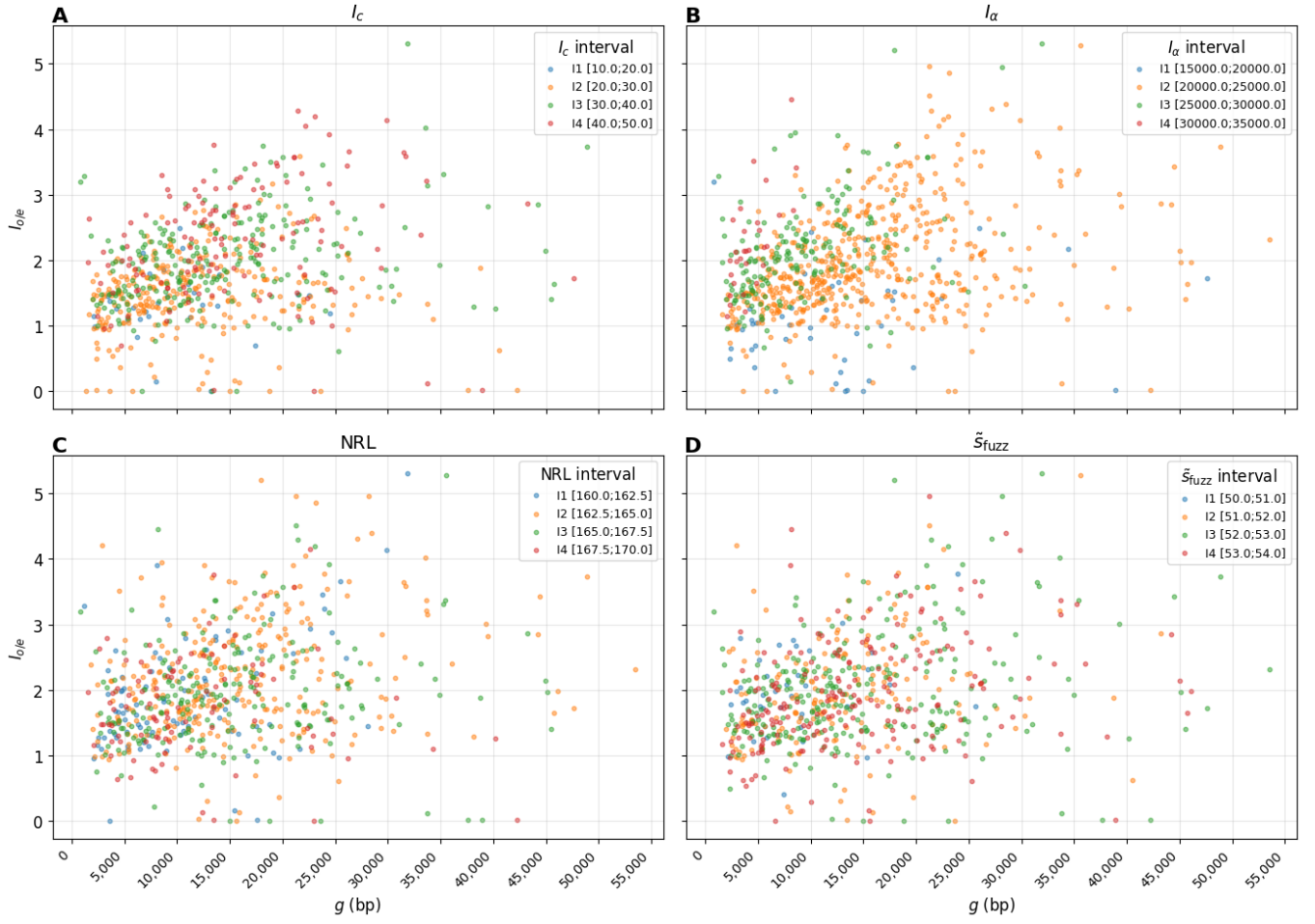

**Figure S15: Correlation between the contact enrichment between consecutive CARs and the properties of the underlying nucleosomal landscape.** For each chromatin properties [$I_c$ (A), $I_\alpha$ (B), $\widetilde{NRL}$ (C), $\tilde{s}_{fuzz}$ (D)], four equal-length intervals are defined covering the full range of values. For all pairs of consecutive CARs from a given interval, we plot the property of interest vs genomic distances $g$ between the CARs.

- [1] Wolfram Research, Inc., Mathematica, Version 14.2, champaign, IL, 2024.
- [2] N. M. Temme, *Special Functions: An Introduction to the Classical Functions of Mathematical Physics* (Wiley, New York, 1996).
- [3] NIST Digital Library of Mathematical Functions, 8.11 asymptotic approximations and expansions (2023), release 1.1.12; accessed 19 February 2026.
- [4] C. Godrèche and J. Luck, *Journal of Statistical Physics* **104**, 489 (2001).
